# Circular *ANRIL* isoforms switch from repressors to activators of *p15/CDKN2B* expression during RAF1 oncogene-induced senescence

**DOI:** 10.1101/2020.04.28.065888

**Authors:** Lisa Muniz, Sandra Lazorthes, Maxime Delmas, Julien Ouvrard, Marion Aguirrebengoa, Didier Trouche, Estelle Nicolas

## Abstract

Long non-coding RNAs (ncRNAs) are major regulators of gene expression and cell fate. The *INK4* locus encodes the tumour suppressor proteins p15^INK4b^, p16^INK4a^ and p14^ARF^ required for cell cycle arrest and whose expression increases during senescence. *ANRIL* is a ncRNA antisense to the *p15* gene. In proliferative cells, *ANRIL* prevents senescence by repressing *INK4* genes through the recruitment of Polycomb-group proteins. In models of replicative and RASval12 oncogene-induced senescence (OIS), the expression of *ANRIL* and Polycomb proteins decreases, thus allowing *INK4* derepression. Here, we found in a model of RAF1 OIS that *ANRIL* expression rather increases, due in particular to an increased stability. This led us to search for circular *ANRIL* isoforms, as circular RNAs are rather stable species. We found that the expression of two circular *ANRIL* increases in several OIS models (RAF1, MEK1 and BRAF). In proliferative cells, they repress *p15* expression, while in RAF1 OIS, they promote full induction of *p15, p16* and *p14*^*ARF*^ expression. Further analysis of one of these circular *ANRIL* shows that it interacts with Polycomb proteins and decreases EZH2 Polycomb protein localization and H3K27me3 at the *p15* and *p16* promoters, respectively. We propose that changes in the ratio between Polycomb proteins and circular *ANRIL* isoforms allow these isoforms to switch from repressors of *p15* gene to activators of all *INK4* genes in RAF1 OIS. Our data reveal that regulation of *ANRIL* expression depends on the senescence inducer and underline the importance of circular *ANRIL* in the regulation of *INK4* gene expression and senescence.

## Introduction

Long non-coding RNAs (lncRNAs) play an important role in the control of gene expression. They are usually involved in the local targeting of chromatin modifying enzymes [1]. Although there are many examples of lncRNAs functioning *in trans*, they usually regulate genes *in cis*, meaning close to where they are produced. One of the best described lncRNA which targets chromatin modifying enzymes to chromatin is the *ANRIL* ncRNA [2]. *ANRIL* is produced from the tumor suppressor *INK4* locus, which contains two protein-coding genes, *CDKN2A* and *CDKN2B*. The *CDKN2B* gene encodes the p15^INK4b^ (p15) Cyclin Dependant Kinase (CDK) inhibitor. The *CDKN2A* gene encodes two proteins, the p16^INK4a^ (p16) CDK inhibitor and p14^ARF^, by a different reading frame using an alternate first exon located 20 kb upstream [3]. *ANRIL* is actually transcribed antisense to the *p15* / *CDKN2B* gene [4]. *ANRIL* has been extensively studied given that it has been identified as a risk locus in many human diseases from Single Nucleotide Polymorphism (SNP) association by Genome Wide Association Studies (GWAS) [5, 6]. Notably, a 53 kb region associated with increased risk of coronary artery diseases (CAD) is located in the second half of the *ANRIL* gene [5]. Increased or reduced *ANRIL* expression has been reported in diverse diseases including cancers, diabetes or cardiovascular diseases, whose risk is linked with aging [7]. Notably, *ANRIL* was found to be overexpressed in many cancers and to promote cell proliferation of cancer cell lines from diverse origins including colon carcinoma [8, 9], esophageal carcinoma and bladder cancers [10, 11]. Moreover, *ANRIL* is a strong candidate gene for being the gene responsible for genetic susceptibility to many inflammatory diseases [12].

The *INK4* locus plays a major role in the control of cellular senescence [13], which is characterized by a stable proliferation arrest. Indeed, the two major pathways that mediate senescence commitment are the p53 and Rb tumor suppressor pathways, which are both regulated by genes encoded by the *INK4* locus [14]. Upon senescence induction, *INK4* gene expression is activated: the p15 and p16 CDK inhibitors activate the Rb pathway by inhibiting its phosphorylation by Cyclin/CDK while p14^ARF^ represses HDM2 function, allowing activation of the p53-p21 pathway. Both events result in a strong cell cycle arrest, which is made irreversible by subsequent events.

Senescence was first described as a stable cell proliferation arrest after a finite number of cell divisions of normal cells in culture (replicative senescence) triggered by telomere shortening in the absence of telomerase activity and the incomplete replication of the very end of chromosomes [15]. It is now clear that senescence is a cell fate induced upon various kinds of stresses, such as oncogene activation, and characterized by a potent and permanent cell proliferation arrest, as well as a number of associated changes, such as chromatin reorganisation and the setting up of a specific genetic program [16]. Senescence is believed to be a major tumour suppressor pathway, through the elimination of cells with oncogenic activation, not repairable DNA damages or uncontrolled genetic instability [17].

*ANRIL* has been shown to prevent senescence induction by repressing the expression of the *p15/CDKN2B* and *p16*/*CDKN2A* genes at the *INK4* locus in proliferative cells by locally recruiting the repressive Polycomb complexes [18, 19]. Polycomb group proteins are involved in transcriptional gene silencing and heterochromatin formation [20, 21]. They are mainly contained within two multimolecular complexes: PRC2 contains the EZH2 histone methyl transferase, which methylates K27 of histone H3. PRC1 specifically recognises methylated H3K27 and mediates transcription inhibition, at least in part through histone H2AK119 monoubiquitinylation [22]. In proliferative cells, *ANRIL* is able to recruit these two complexes *in cis* to the *INK4* locus, allowing its transcriptional silencing [18, 19]. Interestingly, *ANRIL* has also been shown to regulate gene expression *in trans*, although the molecular mechanisms involved are still not clear [23, 24].

During replicative and oncogene-induced senescence (OIS), ChIP experiments indicate that Polycomb group proteins recruitment as well as H3K27 methylation at the *INK4* locus decrease [25-28]. Interestingly, although other mechanisms, such as the general decrease of Polycomb protein expression [25-29], have been described, a senescence-associated inhibition of *ANRIL* expression was proposed to be involved in this process. Indeed, *ANRIL* expression was found to decrease in MEFs undergoing replicative senescence [19] and in a model of OIS induced by oncogenic RAS [18].

Circular RNAs (circRNAs) are ncRNAs produced by back-splicing, which by the splicing of a downstream donor splice site to an upstream acceptor splice site leads to the production of a circular RNA molecule [30]. Since they are naturally resistant to degradation by exonucleases, circRNAs are more stable than linear RNA molecules [31, 32]. Many exonic circular RNAs were found to be mainly cytoplasmic [32, 33] whereas intronic circRNAs are nuclear [34]. CircRNA expression is upregulated during cellular differentiation while their parent gene expression is mostly unchanged [35-38]. CircRNAs are also misregulated in cancers [39]. Recently emerging functions of circular RNAs have been described, especially linked to the regulation of transcription, splicing, protein interaction dynamics or microRNA sponging [40]. Interestingly, many different isoforms of *ANRIL* have been identified [23] including circular *ANRIL* isoforms [41-43] whose expression correlates with anti-proliferative properties and CAD protection [42].

In a recent study, we performed genome-wide analyses of strand specific RNA expression in WI38 human primary fibroblasts undergoing OIS upon activation of RAF1 oncogene [44]. In this experimental setting, *INK4* genes are activated, and their activation is known to be important for the associated cell cycle arrest as well as other senescence features [45-47]. We observed in this model of OIS and other models of OIS that *ANRIL* expression is activated, indicating that repression of *ANRIL* expression is not general during senescence induction and is not required for activation of *INK4* genes. Importantly, we identified several different circular *ANRIL* species whose expression increases in RAF1 induced senescence. Surprisingly, we found that whereas these circular *ANRIL* isoforms repress *p15* gene expression in proliferative cells, they are important for full induction of *INK4* genes in RAF1 induced senescence. Our results indicate that *ANRIL* switches from a negative role on *p15* gene expression to a positive role on *INK4* gene expression during RAF1-induced senescence progression.

## Results

### *ANRIL* expression increases in a model of senescence induced by oncogenic RAF1

In an effort to characterize transcriptional changes in cells undergoing oncogene-induced senescence, we have recently performed strand specific RNA-seq experiments using either proliferative WI38 hTERT RAF1-ER cells (derived from normal human fibroblasts and immortalized by ectopic hTERT expression) or the same cells made senescent by the activation of the RAF1 oncogene [44]. This cell line contains a RAF1-ER (estrogen receptor) fusion protein that allows senescence induction following the addition of 4-hydroxy- tamoxifen (4-HT) [45]. In this *in vitro* model of senescence induction, all cells have stopped dividing after three days of 4-HT treatment and present senescence characteristics such as SAHF formation or the induction of anti-proliferative genes including the *INK4* genes [45, 46]. Close examination of the RNA-seq data at the *INK4* locus, which encodes three major mediators of senescence (p15, p16 and p14^ARF^), indicates that, as previously shown by RT- qPCR experiments [46], *p15, p16* and *p14*^*ARF*^-mRNAs are strongly induced in senescence, with *p15* being particularly activated (Fig. S1A). We also observed that expression of *ANRIL*, which is antisense to the p15-encoding gene and involved in its repression in proliferative cells [18, 19, 48], increases upon senescence induction (Fig. 1A-B and S1B). This observation is surprising given that in the generally accepted model, *ANRIL* expression decreases in senescent cells, resulting in the activation of genes from the *INK4* locus [18, 19].

**Figure 1:**
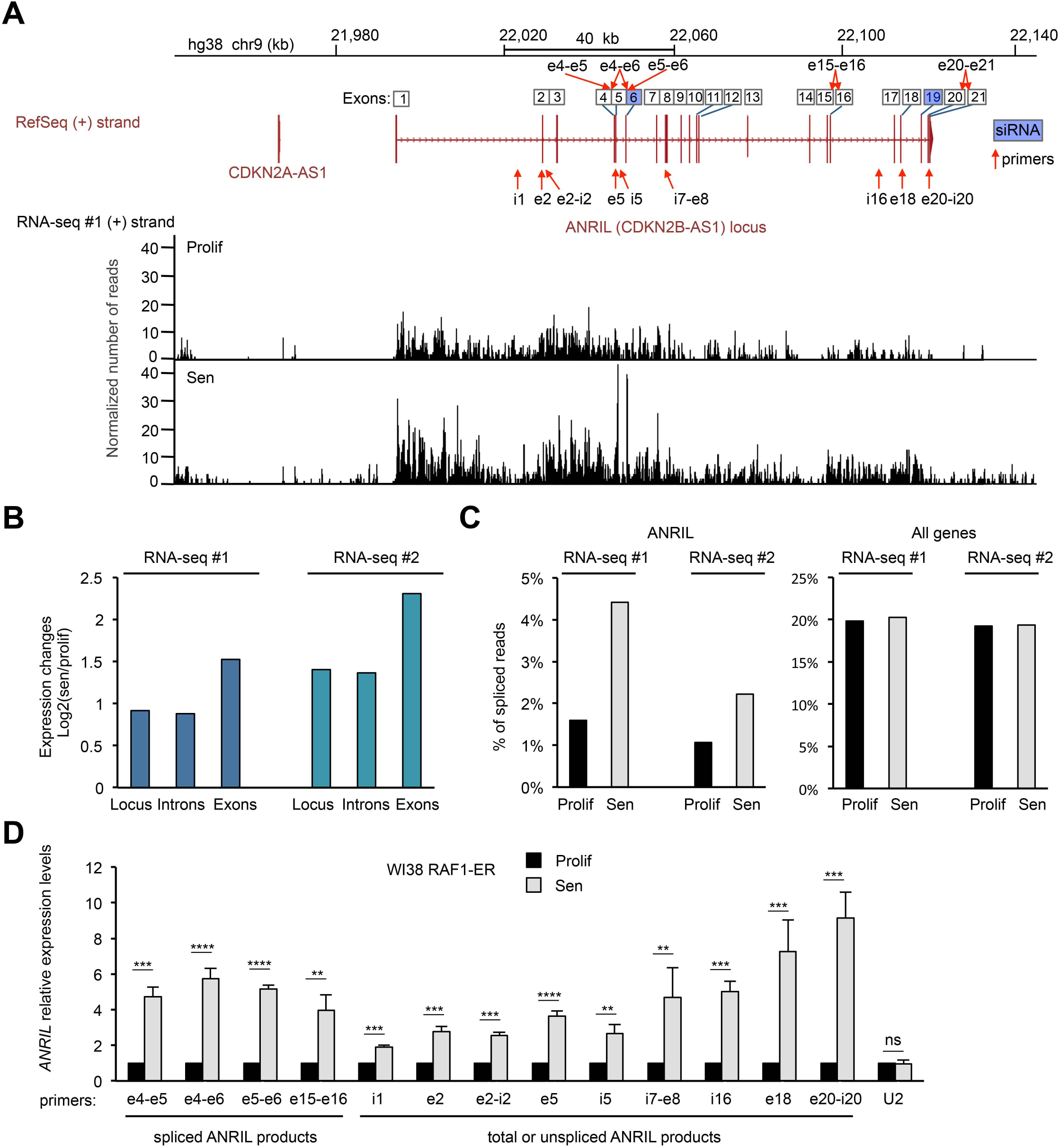
*ANRIL* expression increases in a model of RAF1 oncogene-induced senescence. **A)** RNA-seq data (from the 1^st^ replicate (RNA-seq#1)) showing expression of the (+) strand at the *ANRIL* gene, in WI38 hTERT RAF1-ER cells either proliferative (Prolif) or induced to senescence (Sen) by 4-HT treatment, as indicated. Strand specific RNA-seq tracks show the number of reads per base normalized by the total number of aligned reads multiplied by 100 millions. The *ANRIL* locus is also shown with all the annotated exons from the various transcript variants from the RefSeq database. The primers and siRNAs targeting *ANRIL*, used in this study, are shown with red arrows and blue squares, respectively. Primers detecting spliced products are shown on top of the exon numbers while primers detecting total (spliced + unspliced) or unspliced *ANRIL* are shown below the locus. **B)** Quantification of the RNA-seq signal from the 1^st^ (RNA-seq #1) and 2^nd^ (RNA-seq #2) replicates at the *ANRIL* locus, exons or introns. For each analysed region the mean per base of the normalized number of aligned reads was computed. The ratio of this number in senescence to the number in proliferation was then calculated (log2). **C)** Analysis of strand specific RNA-seq datasets for spliced reads relative to the total number of reads in proliferation or senescence for the *ANRIL* gene or all expressed genes (15,027 genes analysed as described in the Methods section, showing the median value of the whole population). **D)** WI38 hTERT RAF1-ER cells were induced to senescence or maintained proliferative, as indicated. 72 hours later, total RNA was extracted, and *ANRIL* expression was measured by RT-qPCR using the indicated primers. *U2* ncRNA was measured as a control whose expression does not changed in RAF1-induced senescence. The means and standard deviations from 4 independent experiments are shown, relative to *GAPDH* and normalized to 1 in proliferative cells for each experiment (to compare the fold change of *ANRIL* expression during RAF1-induced senescence at different regions. we normalized data to 1 in proliferative cells for each experiment before calculating the mean between different independent experiments). Significant differences are indicated with asterisks (*: p value < 0.05, ** to **** p values < 10^−2^, 10^−3^ and 10^−4^ respectively, two-sided paired Student’s t-test on log2 values). i=introns, e=exons, ns= not significant.

Strikingly, close examination of RNA-seq data shows that the expression of exons seems to increase more than the expression of introns, suggesting that the percentage of spliced forms of *ANRIL* increases in senescence (Fig. 1B). As *ANRIL* has many transcript variants annotated in the reference genome (RefSeq hg38), for simplicity we numbered the exons based on the position of all the exons of these different variants and used this nomenclature throughout this study to name primers and siRNAs (*ANRIL* locus, Fig. S2). We therefore calculated from our two RNA-seq replicates the percentage of spliced reads compared to the total number of reads at the *ANRIL* gene. We found that this percentage is very low in proliferative cells (about 1%), whereas it increases by more than 2 fold in senescent cells (Fig. 1C). This increase is very specific to *ANRIL* since the percentage of spliced reads considering all genes is stable during senescence induction (Fig. 1C). Thus, these data suggest that spliced mature *ANRIL* transcripts specifically increase in senescence. By analysing in details the *ANRIL* spliced junctions found in the RNA-seq experiments, we found that the most abundant splicing event of *ANRIL* in senescence occurs between the exon 5 and the exon 6 (Table S1). Note that small *ANRIL* isoforms, ending at the exon 13, are described in the reference genome (Fig. S2). However, we found only one read containing a spliced junction with exon 13 in only one RNA-seq replicate (data not shown). In contrast, many reads were found connecting the downstream exons in both RNA-seq replicates, suggesting that the small *ANRIL* isoforms are largely underrepresented as compared to large *ANRIL* isoforms ending at exon 21 in WI38 cell line (Table S1). We also observed that *ANRIL* expression increases more in its second part than in its first part (Fig. S3A, B and C).

In order to validate these results, we induced WI38 hTERT RAF1-ER cells to senesce or not. Senescence induction was checked by the analysis of EdU incorporation and SAHF formation (Fig. S4). We first confirmed the increase of *ANRIL* expression at the individual cell level by RNA FISH (Fig. S5). We then recovered total RNAs and confirmed the increase of *ANRIL* expression at the cell population level by Reverse Transcription followed by qPCR (RT- qPCR) at different locations across *ANRIL* using primers detecting only the spliced *ANRIL* (in exon-exon junctions), the unspliced *ANRIL* (within introns), or both spliced and unspliced *ANRIL* referred to as total *ANRIL* (within exons) (Fig. 1D). We also found that *ANRIL* spliced isoforms (measured in e4-e5, e4-e6, e5-e6, e15-e16) increase more than unspliced or total *ANRIL* measured in the first part of the gene (i1, e2, e2-i2, e5 and i5) (Fig. 1D). Moreover, confirming the RNA-seq analysis in Fig. S3, the second half of unspliced or total *ANRIL* transcripts (i7-e8, i16, e18 and e20-i20) increases more than its first part (i1, e2, e2-i2, e5 and i5) (Fig. 1D).

Thus, taken together, these data indicate that in WI38 cells undergoing RAF1-induced senescence, *ANRIL* expression increases during senescence progression, and shows a complex regulation among its transcript variants.

### *ANRIL* stability increases in RAF1-induced senescence

To gain insight into the mechanisms underlying the increase in *ANRIL*’s expression in RAF1- induced senescence, we monitored transcription by capture of nascent transcripts. We found that, as expected, *p15* and *p16* transcription increases in senescence (Fig. 2A). This increase in transcription is comparable to the increase of their steady-state mRNA levels. In contrast, *ANRIL* transcription increased weakly (by 2 fold at most) (Fig. 2A), whereas in the same experiments *ANRIL* RNA expression increased 4 fold (see total RNA levels as compared to nascent RNA levels, measured with primers in exon 5 detecting total *ANRIL*). This result indicates that *ANRIL* transcription is only weakly affected by senescence progression, suggesting that its stability might also be affected. We thus monitored *ANRIL* half-life in proliferative and senescent cells by assessing the decrease in its levels following transcription inhibition [49-51]. As controls, we found that the half-lives of *p15* or *p16* mRNAs do not change much during senescence induction (Fig. 2B). In contrast, *ANRIL* stability is increased, with its half-life of about two hours in proliferative cells (as previously described in a genome wide study of RNA half-lives [52]) increasing to up to more than five hours in senescent cells. Thus, Fig. 2 data indicate that the increase in *ANRIL* expression in WI38 cells undergoing RAF1 oncogene-induced senescence is due to both an increase in its transcription and an increase in its stability.

**Figure 2:**
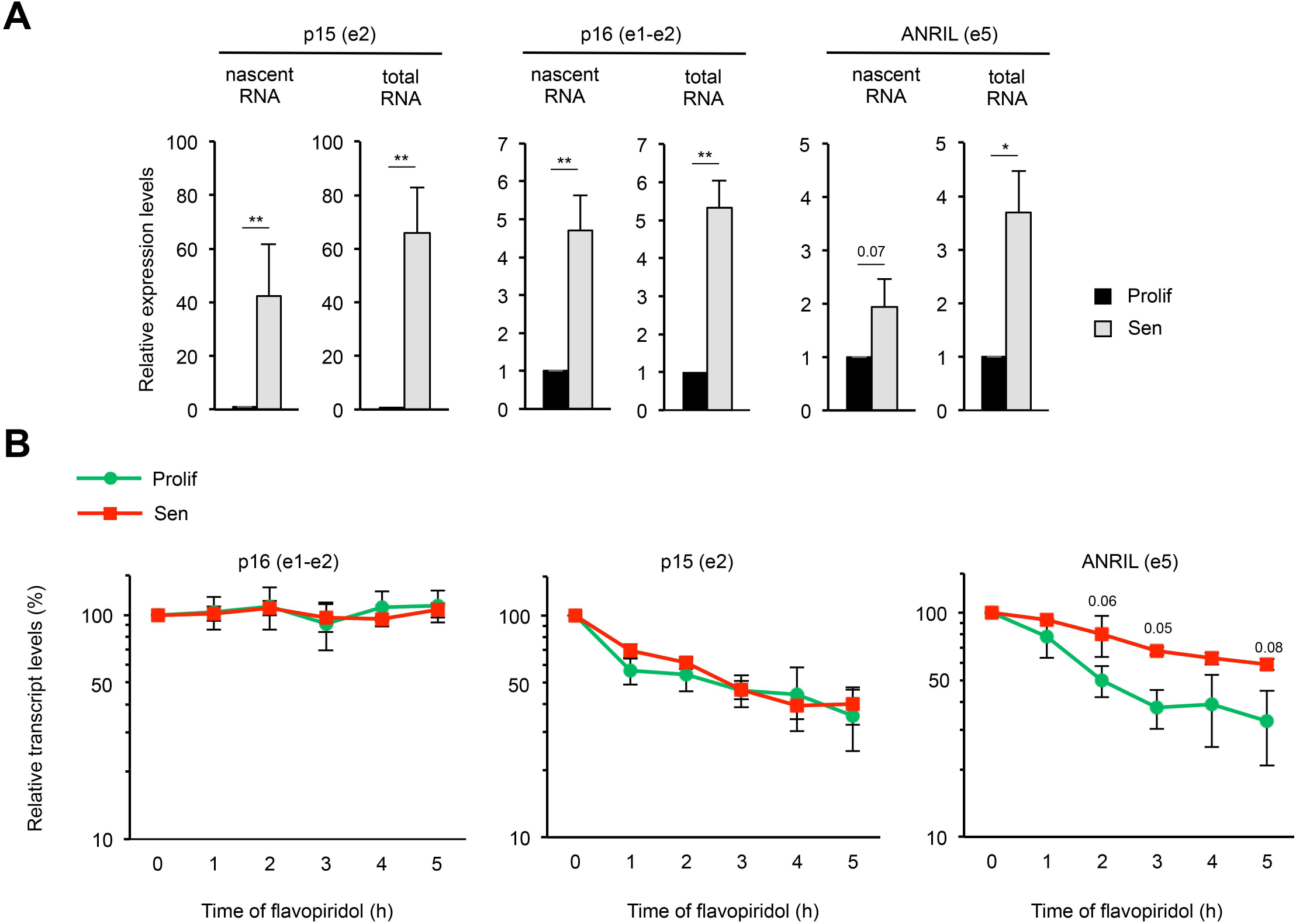
*ANRIL* is stabilized in senescence. **A)** WI38 hTERT RAF1-ER cells were induced to senescence or maintained proliferative, as indicated. 72 hours later, cells were treated with EU for 1 hour, RNA was extracted and EU labelled nascent RNA was purified using the Click-iT technology. Levels of total and nascent *p15, p16* and *ANRIL* were measured by RT-qPCR using the indicated primers. The means and standard deviations from three independent experiments are shown, relative to *GAPDH* (e9) and normalized to 1 in proliferative cells for each experiment. Significant differences are indicated with asterisks (*: p value < 0.05, ** to **** p values < 10^−2^, 10^−3^ and 10^−4^ respectively, two-sided paired Student’s t-test on log2 values) or by the number of the p value when it is between 0.05 and 0.1. **B)** WI38 hTERT RAF1-ER cells were induced to senescence or maintained proliferative, as indicated. 72 hours later, cells were treated with flavopiridol to inhibit transcription for the indicated times. *p15, p16* and *ANRIL* expression was monitored by RT-qPCR using the indicated primers at the indicated times following flavopiridol addition. The levels of RNA were normalised to those of *GAPDH* (e9) and then normalised to 100% at 0 time point for each experiment. The means and standard deviations from three independent experiments are shown (logarithmic scale). Only differences with a p-value between 0.05 and 0.1 are indicated, p-values > 0.1 are not indicated (two-sided paired Student’s t-test on log2 values).

### Circular *ANRIL* RNAs are strongly induced in senescence

Interestingly, many circular *ANRIL* isoforms generated by back-splicing of downstream exons to exon 5 were previously identified [41-43]. Since circular RNAs are more stable than linear RNAs [31, 32, 38], we hypothesized that the increase in *ANRIL* stability could be due to an increase in circular *ANRIL* species. To identify circular forms of *ANRIL* in RAF1-induced senescence, we performed random primed RT followed by PCR using outward facing primers in different exons (Fig. S6). We used primers in exon 5 or exon 6, since exon5-exon6 junction was the most abundant junction found in RAF1-induced senescence in our RNA-seq datasets (Table S1). We also tested the detection of circular isoforms with outward facing primers in exon 16 (because it was found in back-spliced junctions with exon 5 from different studies [41-43]) and in exon 1 (a control since the first exon of a transcript cannot be included in circular RNA molecules since it does not possess splicing signals at its 5’ end) (Fig. S6). After running the RT-PCR reactions on an agarose gel, we detected many products using the outward facing primers in exons 5, 6 and 16, likely representing circular *ANRIL* species. To identify these circular species, the PCR products were purified and sequenced. The circular *ANRIL* species that we have identified in these experiments are represented in Fig. S6 and were previously described in the literature [41-43]. We thus detected back-splicing from exon 6 to exon 5, from exon 7 to exon 5 and from exon 16 to exon 5. These data indicate that circular *ANRIL* species produced by back-splicing at exon 5 are readily detected in RAF1- induced senescent WI38 cells.

Moreover, we observed that *circANRIL* e7-e5 and *circANRIL* e16-e5 are strongly induced in senescence as measured by RT-qPCR using a reverse primer in exon 5 associated with a primer spanning the back-spliced junction (Fig. 3A and 3B). Note that although these primers give a unique PCR product (data no shown), they can detect all circular species containing these specific back-spliced junctions, irrespective of the number or identity of spliced exons or introns located between exon 5 and the reversed spliced exon (Fig. 3A). Finally, although we tried using several primer pairs, we have not been able to specifically detect circANRIL e6-e5 back-spliced junction by RT-qPCR (data not shown) and we thus did not pursue analysis of this *circANRIL* isoform.

**Figure 3:**
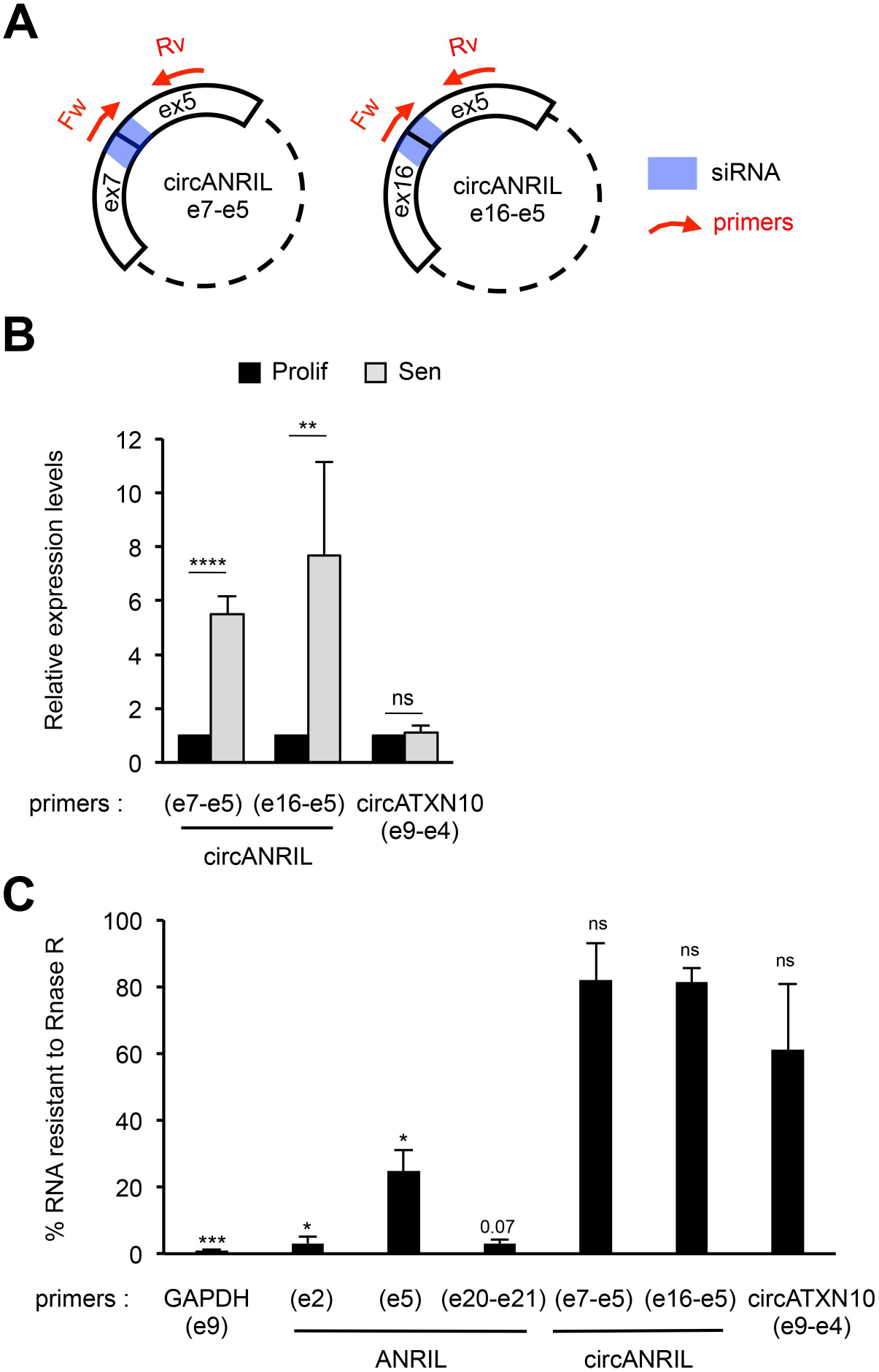
Circular *ANRIL* species are strongly induced in senescent cells. **A)** Schematization of the circular *ANRIL* isoforms containing the specific back-spliced junctions that we identified in our study (Fig. S5). The primers and siRNAs targeting these specific isoforms are shown in red and blue, respectively. **B)** Total RNA was extracted from proliferative or senescent WI38 hTERT RAF1-ER cells. Circular *ANRIL* specie expression was measured by RT-qPCR using the indicated primers. *CircATXN10* was measured as a control whose expression does not changed in RAF1-induced senescence. The levels of RNA were normalised to those of *GAPDH* (e9) and then normalised to 1 in proliferative cells for each experiment. The means and standard deviations from 4 independent experiments are shown. Significant differences are indicated with asterisks (*: p value < 0.05, ** to **** p values < 10^−2^, 10^−3^ and 10^−4^ respectively, two-sided paired Student’s t-test on log2 values), the number of the p value is indicated when it is between 0.05 and 0.1; ns: not significant. **C)** Total RNA from senescent WI38 hTERT RAF1-ER cells were treated or not with RNase R to digest all linear RNAs. Levels of RNAs from treated and untreated samples were analysed by RT-qPCR. Percentages of RNAs resistant to RNase R treatment are calculated relative to the levels of these RNAs measured in untreated samples. Primers designed for e20- e21 junction detect both spliced and unspliced products, the intron 20 being very small (93 nt). Means and standard deviations from 3 independent experiments are shown (only 2 for ANRIL e20-e21, circANRIL e7-e5 and circANRIL e16-e5). Significant differences are indicated as in B.

To gain insights into the proportion of circular *ANRIL* compared to total *ANRIL*, we first analysed splicing in RNA-seq data. We detected 11 reads corresponding to back-spliced junctions in senescence in total, which can be compared to 48 reads containing the e5-e6 junction (present in both linear and circular *ANRIL*) in senescent cells (Table S1), suggesting that about 25% of spliced *ANRIL* are present within circular species in senescent cells. To confirm this result, we made use of RNAse R, which specifically degrades linear RNAs [53]. We found that, as expected, both circular forms of *ANRIL* we analysed above (*circANRIL* e7- e5 and *circANRIL* e16-e5) were largely resistant to RNAse R degradation, whereas virtually all *GAPDH* mRNA was degraded by RNAse R (Fig. 3C). When we analysed exon 5 of *ANRIL*, which is included within all circular *ANRIL* species we identified, we found that about 25% of it was resistant to RNAse R degradation in senescent cells. In contrast, *ANRIL* analysed in exon 2 and in exon 20-exon 21 regions were much more degraded by RNase R treatment, suggesting that these regions of *ANRIL* are mostly included in linear species. Note that exon 20-exon 21 junction degradation is consistent with the fact that the last exon of a transcript cannot circularize as it does not possess splicing signals at its 3’ end (similarly to the first exon, which does not possess splicing signals at its 5’ end). These data suggest that around 25% of the exon 5 of *ANRIL* is present as circular RNAs in senescent cells. Altogether, these data indicate that significant amounts of circular species of *ANRIL* are produced during RAF1-induced senescence progression of WI38 fibroblasts.

### Oncogenic RAF1, MEK1 or RAS –induced senescence leads to a specific regulation of *ANRIL* expression

Although cell cycle arrest is complete at 3 days after 4-HT addition in this RAF1-induced senescence model [45], we wondered whether the changes in gene expression that we observed remained stable after 3 days of senescence induction. By performing long kinetics of senescence induction up to 15 days, we found that total and circular *ANRIL* expression, *p21* (which encodes another Cyclin/CDK inhibitor, p21) expression and *INK4* gene expression increase up to 6-9 days following 4-HT addition (Fig. 4A). After this time point, the level of *ANRIL, p21* and *INK4* gene expression remains stable up to 15 days of 4-HT treatment. Thus, the maximum expression of *ANRIL* transcripts occurs after 6-9 days of 4-HT treatment, as for *INK4* mRNAs, and is constant up to 15 days, indicating that they are not transiently induced in response to RAF1 oncogenic stress but are rather stably associated with the senescent state.

**Figure 4:**
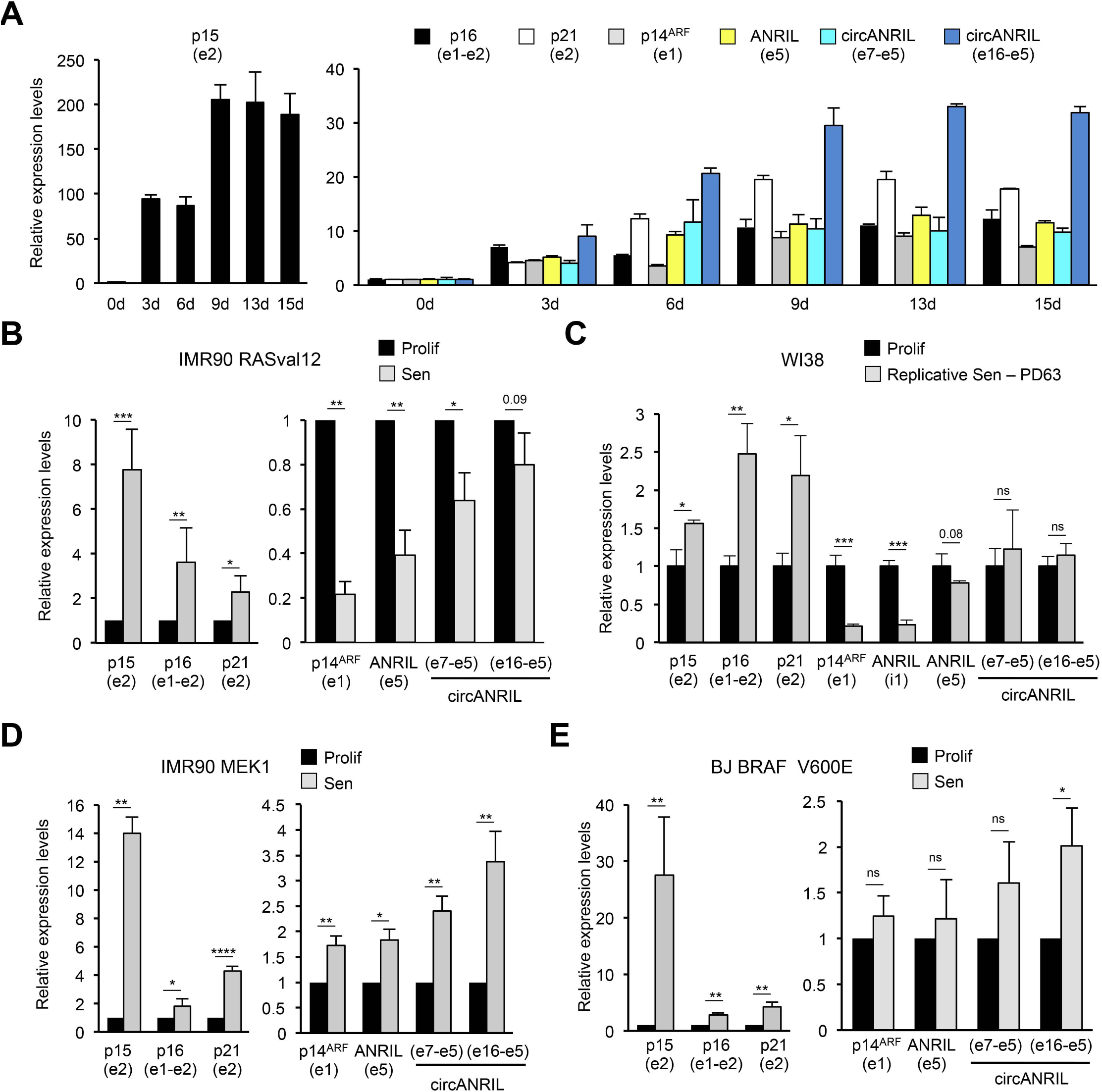
*ANRIL* expression follows *p14*^***ARF***^ **expression in different models of OIS**. **A)** Long kinetics of RAF1-induced senescence. WI38 hTERT RAF1-ER cells were treated with 4-HT for the indicated days (d). RNA expression was analysed by RT-qPCR using the indicated primers. The means and standard deviations from the PCR triplicates of one representative experiment out of 2 is shown, relative to *GAPDH* and normalised to 1 in proliferative cells. **B)** IMR90 RASval12-ER cells were induced to senescence by 4-HT treatment (Sen) or maintained proliferative (Prolif). 8 days later, total RNA was extracted, and RNA expression was measured by RT-qPCR using the indicated primers. The means and standard deviations from 4 independent experiments are shown, relative to *GAPDH* and normalized to 1 in proliferative cells. Significant differences are indicated with asterisks (*: p value < 0.05, ** to **** p values < 10^−2^, 10^−3^ and 10^−4^ respectively, two-sided paired Student’s t-test on log2 values); the number of the p value is indicated when it is between 0.05 and 0.1; ns: not significant. **C)** WI38 cells were grown until replicative senescence (Sen, 63 population doublings (PD)) or arrested while proliferating (Prolif). Total RNA was extracted and RNA expression was measured by RT-qPCR using the indicated primers. The means and standard deviations from 3 independent experiments are shown, relative to *GAPDH* and normalized to 1 in proliferative cells. Significant differences are indicated as in B), except that two-sided unpaired Student’s t-tests were applied. **D)** IMR90 hTERT ΔMEK1-ER cells were induced to senescence by 4-HT addition or maintained proliferative. 3 days after, total RNA was extracted, and RNA expression was measured by RT-qPCR using the indicated primers. The means and standard deviations from 3 independent experiments are shown, relative to *GAPDH* and normalized to 1 in proliferative cells. Significant differences are indicated as in B). **E)** BJ hTERT BRAF V600E cells were induced to senescence by doxycycline addition or maintained proliferative, as indicated. 3 days after, total RNA was extracted, and RNA expression was measured by RT-qPCR using the indicated primers. The means and standard deviations from 3 independent experiments are shown, relative to *GAPDH* and normalized to 1 in proliferative cells. Significant differences are indicated as in B).

*ANRIL* expression was found to decrease in a RASval12 model of oncogene-induced senescence as well as in replicative senescence [18, 19]. This led us to test whether our observation is specific to RAF1-induced senescence of WI38 cells or whether we did not analyse the same *ANRIL* transcript isoforms in our experiments. Using the same primers for RT-qPCR experiments (measuring total *ANRIL* in the exon 5 or *circANRIL* isoforms used in Fig. 1D, Fig. 3B or Fig. 4A), we monitored *ANRIL* expression in an IMR90 cell line immortalized by the ectopic expression of hTERT and expressing an ER-RASval12 fusion allowing, as for RAF1 model, the induction of senescence following 4-HT treatment. Of note, whereas WI38 hTERT RAF1-ER cells were fully senescent in 3 days, IMR90 hTERT ER- RASval12 entered senescence one week after 4-HT addition following one or two cell cycle divisions. We confirmed that, as previously published in WI38 cells induced in senescence by activated H-Ras^G12V^ [18], total *ANRIL* expression decreased about 3 fold in activated RAS-induced senescence (Fig. 4B), in agreement with the generally accepted model. Circular *ANRIL* expression also decreased although to a lesser extent (Fig. 4B). We also monitored *ANRIL* expression in WI38 cells undergoing replicative senescence, and found that, although unspliced *ANRIL* decreases and total *ANRIL* expression slightly decreases, consistently with what has been previously observed [19], circular *ANRIL* levels remain stable (Fig. 4C), Altogether, we conclude from these experiments that *ANRIL* expression is differentially regulated in senescence with respect to senescence inducers.

To investigate whether the increase in *ANRIL* expression is restricted to RAF1-induced senescence, we used other *in vitro* models of senescence. We induced senescence in IMR90 hTERT ΔMEK1-ER cells by treatment with 4-HT for 3 days [54] or in BJ BRAF V600E cells treated with doxycycline for 3 days [55] and monitored expression changes in *ANRIL* and *INK4* genes. In IMR90 hTERT ΔMEK1-ER cells, the expression of total *ANRIL*, circular *ANRIL*, as well as *INK4* and *p21* mRNAs increases with senescence (Fig. 4D). In BJ BRAF V600E cells, whereas total *ANRIL* levels do not change much, the expression of *circANRIL* e16-e5 is significantly induced in senescence. In addition, although it is not significant, the expression of *circANRIL* e7-e5 is induced in all three experiments (Fig. 4E). This indicates that the increase in total or circular *ANRIL* expression is not specific to RAF1-induced senescence nor to WI38 cells.

Strikingly, the evolution of *ANRIL* expression in these various models of senescence parallels that of *p14*^*ARF*^ expression, consistent with the fact that they are produced from divergent promoters. In RAF1-induced senescence, short kinetics of senescence induction indicate that *ANRIL* induction occurs concomitantly with *p14*^*ARF*^ induction or even slightly earlier, and after the induction of *p15* and *p16* (Fig. S7A). Finally, in all the senescence models we tested, we observed that circular *ANRIL* expression increases more or decreases less than total *ANRIL* expression, suggesting that either *ANRIL* circularization is more efficient or that circular *ANRIL* is stabilized in senescent cells.

The long kinetics data shown above as well as its activation in other systems of oncogene-induced senescence suggest that *ANRIL* regulation by RAF1 is linked to the senescence process *per se* and not to RAF1 activation. Accordingly, inhibition of endogenous RAF1 in parental WI38 cells does not affect *ANRIL* production (Fig. S7B), suggesting that *ANRIL* is not directly targeted by RAF1-dependent pathways. To further demonstrate this point, we prevented senescence induction using previously characterized siRNAs directed against p16 and p21 [45]. We found that inhibition of p16 and p21 expression abolishes the activation of both total and circular *ANRIL* expression induced by RAF1 activation in WI38 cells (Fig. S7C), therefore demonstrating that it is linked to the process of senescence.

### *ANRIL* participates to *INK4*-protein coding gene activation in RAF1-induced senescence

We next investigated what could be the role of *ANRIL* and of its circular RNA species in RAF1-induced senescence. We tested whether it could regulate the expression of the gene to which it is an antisense, the p15-encoding gene. To that goal, we used an siRNA-based approach. We first transfected senescent cells with an siRNA targeting exon 6 of *ANRIL*. We found that this siRNA efficiently decreases total *ANRIL* expression measured in exon 5 as well as the expression of both circular *ANRIL* species we analysed above (Fig. 5A). Surprisingly, considering the published role of *ANRIL* in repressing p15-encoding gene expression in proliferative cells, *ANRIL* depletion in senescent cells leads to a decrease in p15-encoding gene expression (Fig. 5A and S8A). Importantly, this was confirmed at the protein level by western blotting (Fig. S8B).

**Figure 5:**
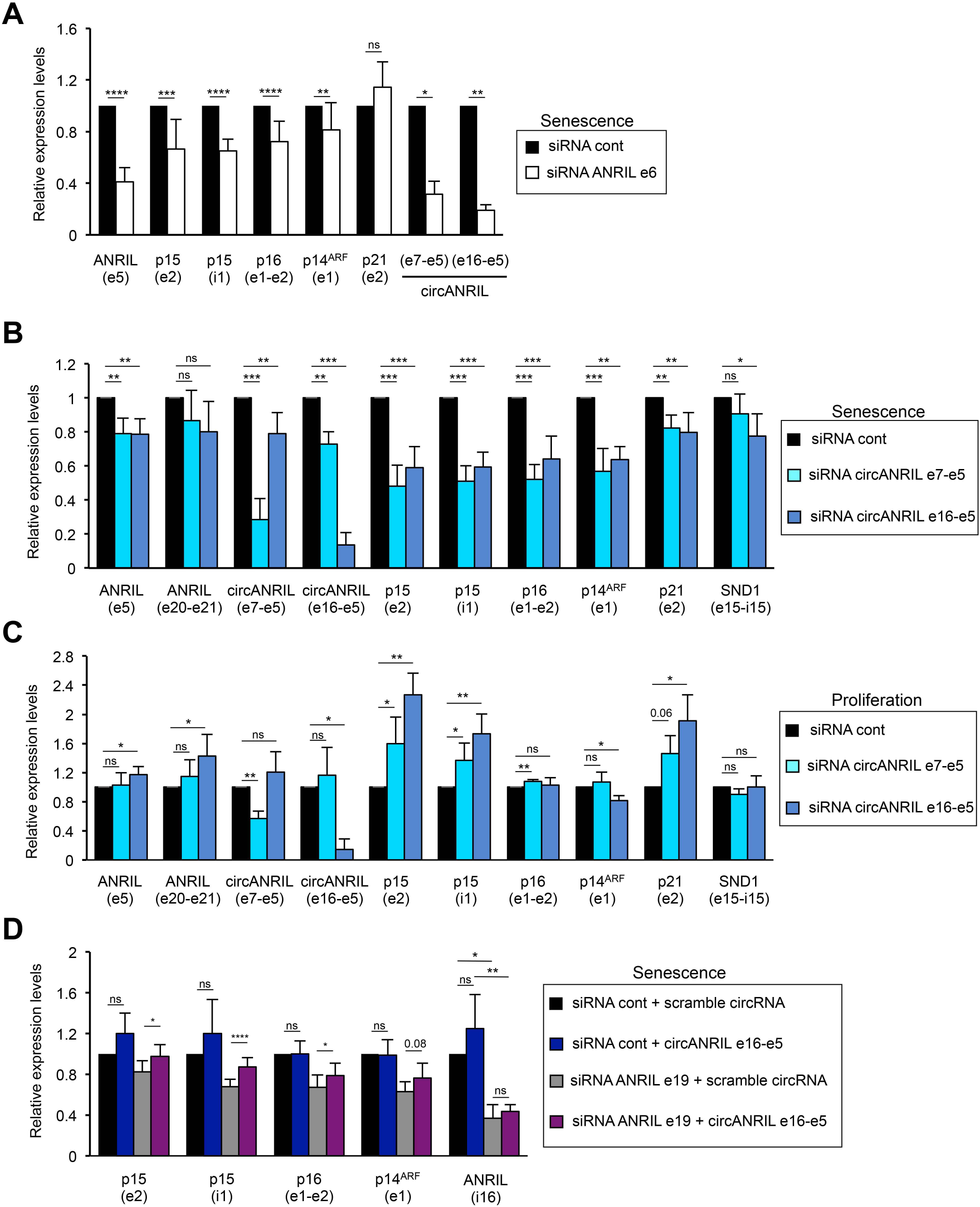
Circular *ANRIL* species participate in *INK4* gene regulation in proliferative or senescent cells. **A)** Senescent WI38 hTERT RAF1-ER cells were transfected with the indicated siRNA. 72h hours later, total RNA was extracted. *ANRIL, p15, p16, p14*^*ARF*^ and *p21* mRNA expression was measured by RT-qPCR using the indicated primers, calculated relative to *GAPDH* and normalized to 1 for the control siRNA in each experiment. The means and standard deviations from 3 to 14 independent experiments are shown, depending on the primers. Significant differences are indicated with asterisks (*: p value < 0.05, ** to **** p values < 10^−2^, 10^−3^ and 10^−4^ respectively, two-sided paired Student’s t-test on log2 values); the number of the p value is indicated when it is between 0.05 and 0.1; ns: not significant. **B)** Same as in A), except that the indicated siRNAs were used. **C)** Proliferative WI38 hTERT RAF1-ER cells were transfected using the indicated siRNAs and analysed as in A). **D)** Senescent WI38 hTERT RAF1-ER cells were transfected with the indicated siRNAs. 24 to 72h hours later, the cells were transfected with the indicated *in vitro* synthesized circular RNAs. 24h later, total RNA was extracted and analysed by RT-qPCR using the indicated primers. The means and standard deviations from 4 independent experiments are shown. Significant differences are indicated as in A).

This result is not due to an off target effect of the siRNA used in these experiments, since it was confirmed using another control siRNA and an siRNA targeting exon 19 of *ANRIL* (Fig. S8C). Note that this siRNA targeting exon 19 of *ANRIL* also decreased circular *ANRIL* expression, although these circular *ANRIL* species do not contain exon 19, indicating that the production of circular *ANRIL* is most likely post-transcriptional.

The p15-encoding gene is located within the *INK4* locus, which also contains the *CDKN2A* gene encoding proteins important for senescence induction: the p16 Cyclin/CDK inhibitor as well as by a distinct promoter, p14^ARF^, an upstream activator of the p53 pathway. We thus analysed the role of *ANRIL* on the expression of the *p16* and *p14*^*ARF*^ mRNAs in senescence. We found that depletion of *ANRIL* led to a decrease in *p16* and *p14*^*ARF*^ mRNA expression but not of *p21* mRNA expression (Fig. 5A and S8C), indicating that in senescent cells, *ANRIL* specifically increases the expression of the senescence-associated genes at the *INK4* locus.

### Circular *ANRIL* species mediate *INK4* gene regulation in proliferative and RAF1 induced-senescent cells

Importantly, siRNAs directed against exon 6 or exon 19 of *ANRIL* strongly affect the expression of the two circular *ANRIL* species we tested (Fig. 5A and S8C). We thus hypothesize that circular *ANRIL* isoforms could play a role in *INK4* gene regulation during RAF1-induced senescence as well as in proliferative cells. We designed siRNAs targeting the back-spliced junction of circular *ANRIL* species to specifically affect their expression (represented in Fig. 3A). siRNAs directed against exon 7-exon 5 or exon 16-exon 5 junctions strongly and specifically affect the expression of their back-spliced products in proliferative and RAF1-induced senescent cells (Fig. 5B-C). They only weakly affect total *ANRIL* measured in exon 5, consistent with the experiments with RNase R from which we estimate that 25% of exon 5 is present within the diverse circular *ANRIL* isoforms (and thus each individual circular specie accounts for a small fraction of total exon 5). Importantly, as expected, they do not affect the expression of linear *ANRIL* species (measured in exon 20- exon 21 junction which, as shown in Fig. 3C, is present within linear species) (Fig. 5B-C).

Depletion of both circular isoforms in RAF1-induced senescence decreased the expression of *p15, p16* and *p14*^*ARF*^ *INK4* gene expression whereas it had only a weak effect, if any, on the expression of *p21* and *SND1* (as a control) genes (Fig. 5B). In RAF1-induced senescence, similar results were obtained when using another siRNA against *circANRIL* e16-e5 compared to a control scramble siRNA (Fig. S9A). To rule out off target effects, we intended to reverse the effect of siRNA by transfection of *in vitro* produced circular RNAs. Given that *in vitro* transcription using T7 RNA polymerase can introduce untemplated nucleotides at the 5’ and 3’ ends of the synthesised RNA, we decided to produce a linear RNA already containing the e16-e5 backspliced junction to avoid introducing mutations in this specific junction. We thus produced a linear RNA containing the 3’ end of exon 7 followed by the entire exon 16-exon 5-exon 6 sequence followed by the 5’ end of exon 7 (Fig. S9B) so that point mutations eventually introduced through addition of untemplated nucleotides by T7 RNA polymerase are located in the middle of exon 7. Upon ligase treatment, ligation of the two halves of exon 7 (producing the entire exon 7), reconstitutes a circularized RNA containing the exact e16-e5 junction and nearly identical to the isoform we identified in senescent cells (see Fig. S9B-C for details of its preparation). In each experiment, we analysed the RNAs purified from transfected cells and found that circularisation was effective but not very efficient (although the e16-e5 backspliced junction was overexpressed on average 1300 times (Fig. S9B), the circularised product was overexpressed around 19 times (data not shown)) and variable from one experiment to the other so that most of the transfected RNAs was linear. Nevertheless, it has to be noted that even the overexpressed linear RNA contains the e16-e5 backspliced junction within its natural context, and may thus mimic the function of *circANRIL* e16-e5 natural RNA, at least to some extent. Most importantly, transfection of this *in vitro* produced RNA containing the e16-e5 backspliced junction reverses the effect of siRNAs decreasing *circANRIL* e16-e5 in RAF1-induced senescence (Fig. 5D and S8C), ruling out off-target effects of siRNAs.

In proliferative cells, depletion of *circANRIL* e16-e5 and *circANRIL* e7-e5 both increase *p15* expression, while *p16, p14*^*ARF*^ and *SND1* expression is largely unaffected (Fig. 5C). These results are consistent with the known function of *ANRIL* in repressing the *INK4* gene locus in proliferative cells [18, 19], suggesting that circular isoforms of *ANRIL* participate in the repression of at least the *p15* gene in proliferative cells. Unexpectedly, the depletion of the two circular *ANRIL* species also increases the expression of *p21* mRNA, suggesting that they can repress anti-proliferative genes *in trans* (Fig. 5C).

Thus, these data indicate that circular species of *ANRIL* are responsible, at least in part, for the repression of *p15* expression in proliferative WI38 fibroblasts, as well as for the full induction of the expression of all *INK4* genes during RAF1-induced senescence progression. Altogether these data suggest that at least two circular *ANRIL* isoforms shift from being repressors of *p15* expression in proliferative cells to activators of *INK4* genes in RAF1- induced senescent cells.

### *CircANRIL* e16-e5 regulates H3K27 modifications at the *p16* gene promoter in RAF1- induced senescence

We next intended to investigate the mechanism by which circular *ANRIL* species positively regulate *INK4* gene expression in RAF1-induced senescent cells. The observation that total *ANRIL* and *circANRIL* e7-e5 or e16-e5 RNAs activate *p15* expression when monitoring *p15* intron expression suggests that this activation occurs transcriptionally (Fig. 5, S8A, S8C and S9A). To test this possibility, we monitored nascent *p15* and *p16* RNAs and found that their expression is decreased upon total *ANRIL* or *circANRIL* e16-e5 and e7-e5 depletion, indicating that they control transcription of *INK4* genes in RAF1-induced senescence (Fig. S10A). To further confirm that circular *ANRIL* isoforms can control transcription of *INK4* genes in RAF1-induced senescent cells, we performed RNA pol II ChIP experiments. We found that depleting *circANRIL* e16-e5 leads to a decrease in RNA pol II recruitment at the *p15* and *p16* promoters (Figure S10B) (We did not analyse *p14*^*ARF*^ promoter, given that it is also the promoter of *ANRIL*, which could affect the results). Thus, *circANRIL* e16-e5 favours the recruitment of RNA pol II at these two promoters in RAF1-induced senescent cells.

We next analysed the consequences of depleting *circANRIL* e16-e5 in RAF1-induced senescent cells on the presence of chromatin marks known to regulate *INK4* gene expression. Among these, we analysed H3K27 methylation and its counteracting modification, H3K27 acetylation, which have both been involved in *INK4* gene regulation [26, 27]. We did not find any significant effect on H3K27 modifications at the *p15* promoter. These results could be interfered by the fact that the *p15* gene is antisense and thus included within the *ANRIL* gene whose expression is increased in senescence. Strikingly, however, we found that the presence of H3K27ac is decreased whereas the presence of H3K27me3 is increased (Fig. 6A) at the *p16* promoter upon *circANRIL* e16-e5 depletion in RAF1-induced senescent cells. Thus *circANRIL* e16-e5 regulates H3K27 modifications, at least at the *p16* promoter.

**Figure 6:**
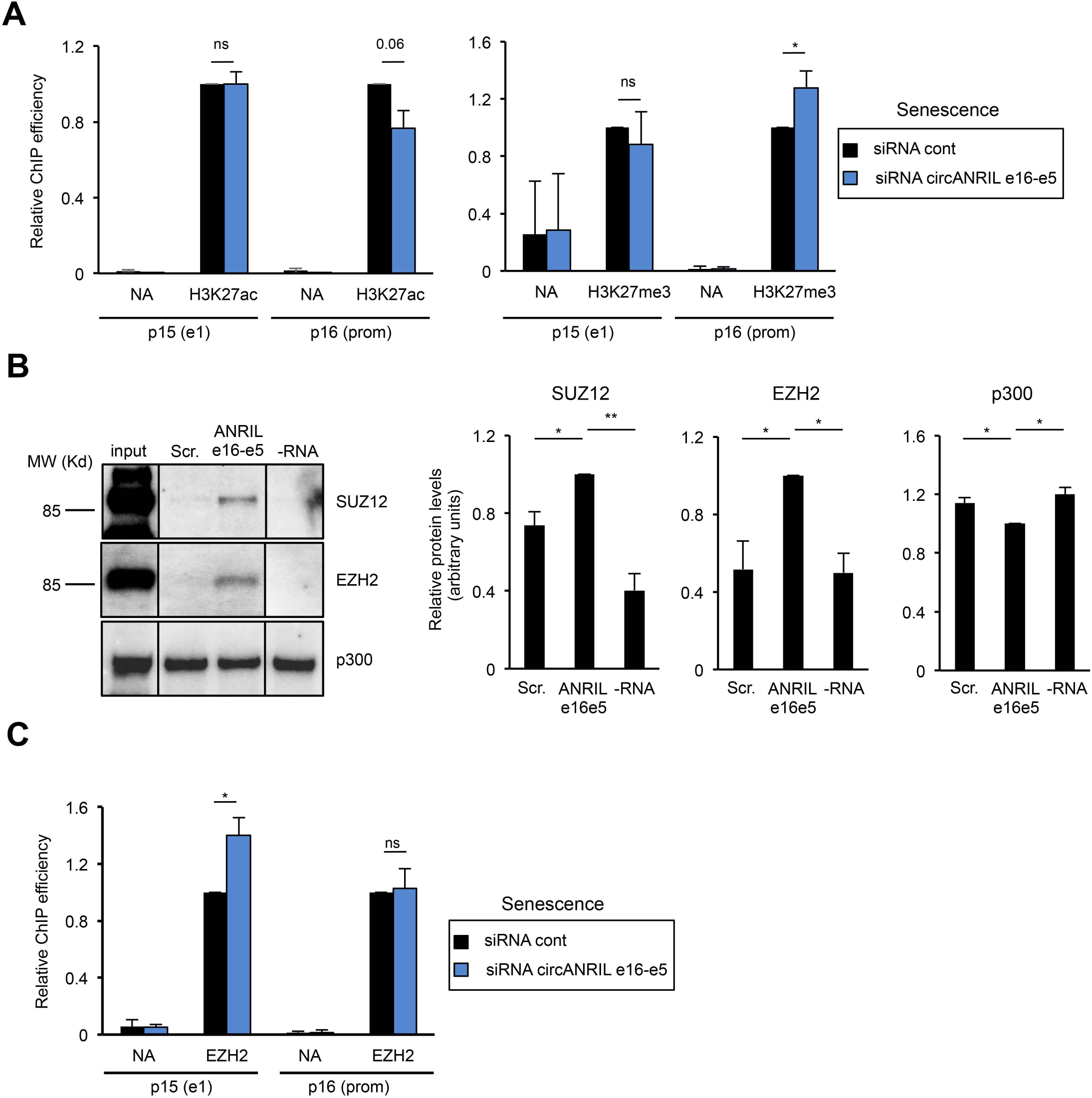
Circular *ANRIL* e16-e5 regulates H3K27me3 at the *p16* promoter in senescent cells. **A)** Senescent WI38 hTERT RAF1-ER cells were transfected with the circANRIL e16-e5 siRNA. Three days later, cells were harvested and subjected to a ChIP analysis using H3K27ac (left panel) or H3K27me3 (right panel) or total histone H3 antibody or no antibody (NA) as a control. The amount of p15 e1, p16 e1 or GAPDH e9 (used as a positive control since it also detects pseudogenes with significant H3K27ac and H3K27me3 levels) sequences were quantified by qPCR. The percentage of input was calculated and the amount of *p15* and *p16* promoters were calculated relative to 1 for the control siRNA sample following standardisation with GAPDH e9 and total nucleosome occupancy (H3 ChIP values). The means and standard deviations from 3 independent experiments are shown. Significant differences are indicated by an asterisk (p value < 0.05, two-sided paired Student’s t-test on log2 values); number of the p value is indicated when it is between 0.05 and 0.1; ns: not significant. **B)** *In vitro* transcribed biotinylated RNAs either containing the ANRIL e16-e5 spliced junction or a scrambled sequence (Scr) were incubated with HeLa nuclear extracts. Biotinylated RNAs were then recovered using streptavidin beads and co-precipitated proteins were analysed by western blotting using SUZ12 or EZH2 antibodies, as indicated. Typical western blots are shown. Lines indicate unnecessary intervening lanes that have been spliced out. The histograms show the means and standard deviations of the means (SDOM) from 8 independent experiments (for SUZ12 and EZH2) or 5 independent experiments (for p300) excluding outliers. Significant differences are indicated with asterisks (*: p value < 0.05, **: p values < 10^−2^, two-sided paired Student’s t-test on log2 values), ns: not significant. **C)** Same as in A), except that an anti-EZH2 antibody was used and only GAPDH e9 but not total nucleosome occupancy was used for standardisation.

To test whether this effect could be direct, and given that *in vitro* produced RNAs containing the *circANRIL* e16-e5 junction can reverse the effect of siRNAs on *INK4* gene expression (see above), we checked whether enzymes regulating H3K27 modifications could interact with it. H3K27 tri-methylation is mediated by EZH2, an histone methyltransferase, which is the catalytic subunit of the PRC2 Polycomb complex, whereas H3K27 acetylation is mediated by CBP/p300 [22]. We incubated *in vitro* produced biotinylated RNAs containing the *circANRIL* e16-e5 junction with nuclear extracts and recovered interacting proteins using streptavidin beads. No specific binding of p300 to *circANRIL* e16-e5 junction-containing RNAs was observed; p300 being pulled-down by streptavidin beads even in the absence of biotinylated RNA (Fig. 6B). In contrast, we found that SUZ12 and EZH2 proteins are more efficiently recovered using RNAs containing the *circANRIL* e16-e5 junction than using control scramble RNAs (Fig. 6B). Moreover, we found that, depending on the experiment, from 1.2 % to 13% of the input EZH2 or SUZ12 were recovered with *in vitro* produced RNAs containing the *circANRIL* e16-e5 junction. Thus, these data indicate that RNAs containing the *ANRIL* e16-e5 backspliced junction can specifically interact with the SUZ12 and EZH2 Polycomb group proteins.

The PRC2 complex containing EZH2 has been shown to be directly recruited to the *INK4* locus [27, 28]. We thus investigated whether *circANRIL* e16-e5 regulates EZH2 recruitment to the *INK4* locus. In RAF1-induced senescence, we found that *circANRIL* e16-e5 depletion increases EZH2 recruitment at the *p15* promoter whereas no effect could be observed at the *p16* promoter (Fig. 6C). These data suggest that *circANRIL* e16-e5 regulates the recruitment of EZH2 to the *p15* promoter of the *INK4* locus during RAF1-induced senescence, resulting in changes of H3K27me3 methylation at the *p16* promoter, probably because of chromatin folding [56].

Altogether, these data indicate that some circular species of *ANRIL* favour the transcription of p15- and p16-encoding genes in RAF1-induced senescence probably by sequestering EZH2 and preventing it from being recruited to some regions of the *INK4* locus, therefore decreasing the presence of the repressive H3K27 trimethylation epigenetic mark.

### *CircANRIL* e16-e5 is important for the genetic program of senescence

Given the importance of *INK4* genes in the control of cell proliferation and senescence induction, we next analysed whether circular *ANRIL* species are important for these processes. In proliferative cells, depletion of total *ANRIL, circANRIL* e16-e5 and e7-e5 leads to a decrease in cell proliferation (Fig. S11), in agreement with the increased expression of *p15* and *p21* we showed above. Depletion of these RNAs in senescent cells, alone or in combination with p16 and/or p21 inhibition does not increase the number of cells able to proliferate (measured by EdU staining) and does not decrease SAHF presence measured by DAPI heterogeneity (data not shown), suggesting that it is not sufficient to revert senescence. Given that *trans* effects of *ANRIL* on gene expression were reported [23], we analysed whether it could be important for the genetic program associated with senescence besides regulation of *INK4* genes. We thus performed RNA-seq experiments following depletion of *circANRIL* e16-e5 in senescence and performed a differential analysis of gene expression. We found 296 genes up-regulated and 294 down-regulated by more than two fold (see Table S2 for the list of regulated genes). A gene ontology analysis on these lists of genes up- or down- regulated upon *circANRIL* e16-e5 depletion can be found in Tables S3 and S4, respectively. Of note, with respect to senescence features, genes induced upon *circANRIL* e16-e5 depletion are enriched in inflammation-linked genes and include three genes from the SASP (genes coding for IL6, IL15RA and FGF), whereas genes down-regulated are linked to regulation of cell proliferation.

To test whether *circANRIL* e16-e5 could be important for the genetic program associated with senescence, we crossed these lists of *circANRIL* e16-e5-regulated genes with genes differentially regulated upon senescence induction in the same cells [44]. Strikingly, amongst the 296 genes repressed by *circANRIL* e16-e5 (upregulated upon *circANRIL* e16-e5 depletion), 115 are also repressed in senescence. This is much more than expected by chance, a difference which is highly significant (53 expected, p-value 1.85*10^−21^, chi2 test). Similarly, 78 genes activated by *circANRIL* e16-e5 (downregulated upon *circANRIL* e16-e5 depletion) are also activated in senescence, again significantly different from what expected by chance (26 expected, p-value: 2.81*10^−27^). These data thus indicate that *circANRIL* e16-e5 is important for the genetic program associated with RAF1-induced senescence.

## Discussion

Here, we show that, contrary to the proposed model based on the analyses of *ANRIL* expression changes in Rasval12-induced and replicative senescence [18, 19], senescence is not always associated with a decrease in the expression of *ANRIL*. Instead, we observe in various models of oncogene-induced senescence that *ANRIL* expression increases during senescence. This indicates that, depending on the inducers of senescence, the cell response at the *INK4* locus is different, at least for *ANRIL* and *p14*^*ARF*^ expression. Indeed, although RAS activation is upstream RAF1 and MEK1 activation in the ERK MAP kinase hyperactivation pathway [57], RAF1-induced senescence differs from RASval12-induced senescence. Indeed, in contrast to RAF1-induced senescence, senescence following RASval12 activation requires replication stress, ROS activation and a proliferation burst before entering senescence [45]. The flattened cell morphology of RAS-induced senescent cells also differs from that of RAF1-induced senescent cells, which adopt a fusiform cell morphology [45]. In addition to the activation of the RAF-MEK-ERK pathway, RAS activation was found to activate other pathways, which could explain the differences observed between RAF1- and RASval12- induced senescent phenotypes [57]. In addition to changes in *ANRIL* expression, our data uncover a complex regulation among its transcript variants that we did not fully investigate in this manuscript. Obtaining a complete picture of the whole spectrum of *ANRIL* variant expression changes associated with senescence would require a much more thorough analysis. In this manuscript, we focused on two circular *ANRIL* species, given that the function of circular isoforms of *ANRIL* was not previously investigated during senescence induction.

Our data also indicate that a decrease in *ANRIL* expression is not absolutely required for *p15* induction during senescence. Indeed, *p15* is strongly induced during RAF1-, BRAFV600E- and MEK1-induced senescence, while the decrease in *ANRIL* expression is absent. This indicates that the relief of *p15* silencing by Polycomb complexes could be mediated by other mechanisms than the decrease of *ANRIL* expression, such as the documented decrease of Polycomb group proteins expression [25-29]. Moreover, recent data demonstrate the critical role played by the MSK kinase, which, by phosphorylating H3S28, relieves Polycomb-mediated repression at the *INK4* locus in oncogene-induced senescence [58].

In agreement with the previously reported function of *ANRIL* on *INK4* gene expression in proliferative cells [18, 19], we indeed found that in our cell model circular *ANRIL* represses *p15* expression in proliferative cells. However, we further describe for the first time that *ANRIL* participates in the activation of *INK4* encoding genes during RAF1-induced senescence. Other studies support a role of *ANRIL* in the activation of the expression of the *INK4* locus. As such, inhibiting *ANRIL* expression has been reported to slightly decrease *p15* expression in colon cancer cell line [8]. Moreover, deletion of the mouse region orthologous to the human chromosome 9p21 CAD risk locus, which is localized in the second half of the *ANRIL* human gene [5], leads to reduced *INK4* gene expression [59].

Thus, the fact that *ANRIL* can participate in the full activation of *INK4* gene expression [8, 59], as we show here during RAF1-induced senescence, could provide the basis for data showing a positive correlation between *ANRIL* and protein-coding *INK4* gene expression in normal and pathological tissues [4, 41, 60-66].

Here, we also uncover that circular *ANRIL* expression can be regulated during a physiological cell response, i.e. senescence induced by the activation of oncogenes. Indeed, we identified two circular *ANRIL* species, circ*ANRIL* e7-e5 and e16-e5, whose expression greatly increases in RAF1-induced senescence as well as in MEK1- or BRAFV600E- induced senescence (Fig. 3 and 4). This increase in circular *ANRIL* isoforms, at least in the RAF1-induced model of senescence, likely explains the increase of total *ANRIL* stability (measured in exon 5) (Fig.2B). RNA circularization can be regulated by the binding of specific factors to linear RNAs [35, 67], but also by the rate of transcription elongation [38]. *ANRIL* circularization could thus be differentially controlled between proliferative and senescent cells by specific interactions with back-splicing regulators. However, it is tempting to speculate that increased production of circular *ANRIL* species is due to a change in the transcription elongation rate at the *ANRIL* locus during RAF1-induced senescence. Indeed, we have previously shown that transcription elongation rate is increased at some loci in RAF1-induced senescence [44].

Most importantly, we demonstrate that these circular isoforms of *ANRIL* play an important function and can actually recapitulate *ANRIL* functions on *INK4* gene expression regulation as previously shown for *ANRIL* in proliferative cells [18, 19] and in RAF1-induced senescence (this study). Indeed, we found that these circular forms of *ANRIL* repress p15-encoding gene expression in proliferative cells and activate *INK4* gene expression in RAF1-induced senescent cells (Fig. 5). Such a role of circ*ANRIL* isoforms in RAF1-induced senescence could also be shared in other models of OIS such as in MEK1- and BRAFV600E in which we show that the expression of circ*ANRIL* isoforms also increases. Note that we were not able to demonstrate whether linear *ANRIL* species can also mediate *ANRIL* function. Indeed, siRNAs targeting the first or the last spliced junction, which cannot be included in circular species were inefficient (data not shown). Moreover, the formation of circular RNAs is largely post-transcriptional [38] and can occur after alternative splicing of their precursor linear RNAs [40, 68]. Our results showing that targeting the exon 19, which is not included in the circular species identified here, decreases the level of all circular *ANRIL* species we tested (Fig. S8C), indeed indicate that *ANRIL* circularization occurs post-transcriptionally. As such, we were unable to specifically inactivate linear RNAs without affecting the circular isoforms since linear RNAs are actually precursors of the circular isoforms.

What could be the mechanisms by which circular *ANRIL* isoforms regulate *INK4* gene transcription and switch from being repressors of *p15* to activators of all *INK4* genes during RAF1-induced senescence progression?

In proliferative cells, circ*ANRIL* isoforms could participate, together with other species of *ANRIL* [18], in the recruitment of Polycomb proteins at its site of transcription to repress the expression of the p15-encoding gene. Indeed, *circANRIL* e16-e5 can interact with Polycomb proteins (Fig. 6B). In proliferative cells, this mechanism seems to be restricted to the p15- encoding gene amongst *INK4* genes, since p16- and p14^ARF^-encoding genes are not regulated by *circANRIL* species (Fig. 5C). This specificity could be related to the fact that *ANRIL* is antisense to *p15* but not to the other *INK4* genes and is consistent with previous observations for total *ANRIL* [18].

In RAF1-induced senescence, transfection of *in vitro* produced *circANRIL* e16-e5 reverse the effect of siRNAs suggesting that *circANRIL* e16-e5 can function in *trans* to regulate *INK4* gene expression. Moreover, we found that it inhibits the binding of EZH2 to the *p15* promoter. This suggests that, through its interaction with the PRC2 complex, *circANRIL* e16- e5 competes with a factor important for recruitment of PRC2 at this target site. It is tempting to speculate that this factor is linear *ANRIL* isoforms localized at their transcription site. Indeed, linear *ANRIL* isoforms carrying the e1-e2 junction have been shown to participate in PRC2 recruitment on the *p15* gene (18), which is located on the antisense strand within the intron 1 of the *ANRIL* gene. On the *p16* promoter, which is outside the *ANRIL* gene and where PRC2 recruitment is not affected by *circANRIL* e16-e5 depletion, H3K27 marks could be regulated by *circANRIL* by chromatin folding between the *p15* and *p16* promoters within the *INK4* locus [56]. Of note, this mechanism could operate on other loci besides the *INK4* locus, in particular on genes down-regulated upon *circANRIL* e16-e5 depletion. Importantly, such a model of competition for binding to an RNA binding protein is similar to what has been proposed in the case of *circANRIL* function on ribosomal RNA maturation in atherosclerosis [42].

How then does *ANRIL* shift from being a repressor to an activator of *INK4* gene expression during senescence progression? We propose that this is linked to the senescence-associated change in the relative ratio of Polycomb proteins and circular *ANRIL* species, resulting from the increase in circular *ANRIL* expression (this study) but also from the very strong decrease of Polycomb protein expression [25-29], including EZH2 [25, 27, 69]. In proliferative cells, the excess of Polycomb relative to *ANRIL* would ensure that all *ANRIL* species present at the *INK4* locus locally recruit Polycomb proteins. Upon senescence induction, an increased amount of circular *ANRIL* species would sequester the residual Polycomb proteins outside of the *INK4* locus and prevent them from being recruited to the *INK4* locus. For example, this could occur while the excess of circular *ANRIL* RNAs is taken in charge by the RNA export machinery to the cytoplasm. Here, we thus propose a mechanism, relying on changes in the ratio between a non-coding RNA and its effector, by which a non-coding RNA can shift from being a negative to a positive regulator of the same genes during commitment into a specific cell fate. Such mechanisms could participate in increasing the difference in the expression of genes crucial for cell fate control, participating in establishing the strong and irreversible changes that are associated with cell fate commitment.

## Materials and Methods

### Cell culture

WI38 hTERT RAF1-ER, IMR90 RASval12-ER and BJ BRAFV600E cells were a kind gift from Dr C. Mann (CEA, France) [45, 55, 70]. IMR90 hTERT MEK1-ER cells [54] were a kind gift from Dr M. Djabali. WI38-, IMR- and BJ- derived cells were grown in MEM supplemented with L-glutamine, non-essential amino acids, sodium pyruvate, penicillin– streptomycin and 10% fetal bovine serum in normoxic (5% O2) culture conditions. For senescence induction in WI38 hTERT RAF1-ER and IMR90 hTERT MEK1-ER, cells were treated with respectively 20nM or 100 nM of 4-HT (Sigma) for 3 days. IMR90 RASval12-ER cells were treated with 100 nM 4-HT for 8 days (cells were dividing once before they completely stopped proliferating). BJ BRAFV600E cells were treated with 1µg/mL doxycycline for 3 days. For replicative senescence, human embryonic fibroblasts WI38 were grown until 49 population doublings for proliferative conditions or until they entered senescence after ∼63 population doublings (cells were harvested 4 weeks after their last division). During long kinetics of RAF1-induced senescence, the medium (including 4-HT) was changed every 3 days. In parallel to senescent cells, proliferative cells were supplemented with the same volume of ethanol used to treat senescent cells with 4-HT. For transcription inhibition, cells were treated with 1µM flavopiridol for the indicated times (Sigma). siRNA transfection was performed using the Dharmafect 4 reagent (Dharmacon) according to the manufacturer’s recommendations, except that 100nM of siRNA were used and then diluted twice 24 hours later by adding the same volume of medium. Cells were harvested 72h following transfection. For the transfection of senescent cells, cells were treated 72h with 4- HT, then transfected and cultured without 4-HT for 72h. For samples treated with control siRNAs (which do not target any sequence in the genome), either one single control siRNA (cont3 or scramble) or a pool of 8 control siRNAs (thus diluting each siRNA control to avoid potential off-target effects of each of them (pool sicont)) were transfected. siRNA sequences are available in the Table S5.

### Chromatin Immunoprecipitations

ChIP was performed as previously described (46) except that Nuclear lysis buffer was diluted twice before use and chromatin samples were diluted 5 times in dilution buffer.

### Antibodies

Antibody to the GAPDH protein was purchased from Millipore (MAB374). Antibodies to the p15 (sc-612) and p300 (sc-585) proteins were purchased from Santa Cruz. Antibodies to SUZ12 (Ab12073), H3 (Ab1791), H3K27Ac (Ab4729) and H3K27me3 (Ab6002) epitopes were purchased from AbCam. The antibody to EZH2 (AC22, #3147) was purchased from Cell Signaling. The antibody to RNA pol II was purchased from Bethyl (A304-405A).

### Analysis of RNA-seq datasets

RNA-seq experiments, normalization and calculation of RNA-seq ratios between senescent and proliferative samples within specific regions were previously described in [44]. For the analysis of spliced reads, the numbers of aligned reads in the *ANRIL* locus or in all gene loci (28,445; from RefSeq database) were calculated. Genes were only analysed if 10 or more reads were aligned on the gene for each sample and each replicate (15,027 genes). For the same regions, the number of aligned reads, which did not align consecutively on the genome, were calculated and referred to as spliced reads. The percentage of spliced reads was calculated by computing the ratio of the number of spliced reads over the number of total aligned reads for each region. The start and the end of the gap for each spliced sequence were analysed for the *ANRIL* locus (Table S1).

For back-spliced reads analysis (Table S1), a new pipeline was developed including three different detection tools in combination (CircExplorer2, CircRNA finder and FindCirc to detect all potential circular RNA reads), and an annotation step of the identified back-spliced junctions (Delmas et al, manuscript in preparation).

For RNA-seq following depletion of *circANRIL* e16-e5, 5-10 µg of total RNA, extracted as described below in (RNA extraction and Reverse transcription), was submitted to EMBL-GeneCore, Heidelberg, Germany. Two replicates of each sample were sequenced. We used strand-specific RNA-seq method, relying on UTP incorporation in the second cDNA strand. RNA-seq samples were sequenced using Illumina NextSeq 500 sequencer, paired-end, 80-bp reads. The quality of each raw sequencing file (fastq) was verified with FastQC [71]. Files were aligned to the reference human genome (hg38) in paired-end mode with STAR Version 2.5.2 and processed (sorting and indexing) with samtools [72]. Raw reads were counted, per gene_id, using HT-seq Version 0.6.1 [73] on the NCBI refseq annotation gtf file from UCSC in a strand specific mode with default parameters. Genes with a mean of less than one read count for all the samples were eliminated. Differential analysis was performed with DESeq2 Bioconductor R package, Version 1.22.1 with default parameters but using normalization by the total read number per sample. Genes of interest were selected when |log2FoldChange| was higher than 1 and adjusted p-value lower than 0.05. For comparison between the genes regulated by *circANRIL* e16-e5 and genes regulated in senescence, proliferation and senescence RNA-seq datasets were processed exactly the same way by HT-seq and DESeq2.

### EdU and Hoechst staining

EdU and Hoechst staining were performed as previously described (44).

### RNA extraction, RT, qPCR and outward facing PCR

Total RNA was extracted from cells using the MasterPure RNA Purification Kit from Epicentre following the manufacturer’s protocol, except that we used 5µL of 50µg/µL proteinase K for cellular lysis. Alternatively, TRIzol was used. After total nucleic acids precipitation, we proceeded to removal of contaminating DNA using a mixture of DNase I and Baseline Zero DNase, supplemented with Riboguard RNAse inhibitor, for 45min at 37°C. Usually, 500ng of total RNA was used for reverse transcription (RT) using the Superscript III Reverse Transcriptase (Invitrogen). A minus RT reaction was performed for each sample and analyzed by qPCR for the expression of *GAPDH* (using GAPDH e9 primers) to verify the non-DNA contamination. qPCR were performed using the SYBR premix Ex Taq from Takara and the Biorad CFX thermocycler. *GAPDH* expression (measured using GAPDH e9 primers) was used for normalisation between samples in each RT-qPCR experiment.

Outward facing PCR reactions were performed on 6ng of random primed cDNA using the GoTaq G2 DNA polymerase (Promega) following the manufacturer’s recommendations in the Techne TC-3000 PCR thermal cycler (initial denaturation of 2 min 95 ^°^C, followed by 35cycles of 30 sec 95 ^°^C, 30 sec 58 ^°^C, 3min 72 ^°^C, final extension of 5 min 72 ^°^C). PCR products were then run on a 1% agarose gel and visualised using GelRed Nucleic Acid Gel Stain (Biotium). Only well separated bands corresponding to the main PCR products obtained were cut out from the gel and PCR products were extracted from gel slices using GenElute Agarose Spin Columns (Sigma-Aldrich). The PCR products were then ethanol precipitated, resuspended in 10µL of water and sent to Eurofins MWG for sequencing using one of the primers used for the PCR reaction.

Sequences of primers are available in the Table S5.

### RNase R treatment

3.5µg total RNA was incubated for 5min at 65°C and immediately placed on ice. Total RNA was then treated for 15min at 37°C with 10U RNase R from Epicentre in a 20 μl reaction volume. A parallel reaction was performed without adding RNase R. 180 μl of TE buffer (10 mM Tris and 1 mM EDTA) was added to each reaction. Purification and precipitation of RNA were performed using the MasterPure RNA Purification Kit from Epicentre by adding 200 μl of 2X T and C Lysis Solution and following the manufacturer’s protocol. 500ng of untreated total RNA from the parallel reaction and an equal volume of the RNase R treated RNA sample were used for random primed Reverse transcription with Superscript III Reverse Transcriptase.

### Labelling and capture of nascent RNA

Nascent RNA was labelled and captured using the Click-iT Nascent RNA Capture Kit from Life Technologies following the manufacturer’s protocol. Cells were incubated with EU (5- Ethynyl Uridine), an analog of uridine, at a concentration of 0.2mM for a 30min or 1hour incubation and of 0.5mM for a 10 min incubation as specified in the figure legend for each experiment. 1µg total RNA was used for the click reaction.

### *In vitro* RNA synthesis for rescue experiments

10µg of plasmid (sequences can be found in Table S5) were digested overnight using PvuII for scramble and SmaI for circANRIL. Linearised plasmids were used as template for *in vitro* transcription using T7 RNA polymerase (Promega) and a ribonucleotide set (NEB) following the manufacturer’s recommendations for 2h at 37°C. *In vitro* transcribed RNA were incubated for 1h at 37°C after addition of 10units DNase I and 10units DNase Zero (Epicentre). The RNAs were then ethanol precipitated and resuspended in water. For two of the rescue experiments (out of 4 independent experiments), the *in vitro* transcribed RNA were directly used for ligation at this step. However, because we found that the ligation efficiency was very low, we modified the protocol to improve this ligation efficiency. For the other rescue experiments, *in vitro* transcribed RNA were then dephosphorylated using 2units Alkaline Phosphatase (FastAP, Thermo Scientific) for 30min at 37°C following the manufacturer’s recommendations. De-phosphorylated RNAs were purified by phenol/chloroform extraction followed by ethanol precipitation, re-phosphorylated using 10units T4 PNK (Promega) for 30min at 37°C and ethanol precipitated. Around 500ng of RNA were ligated overnight at 16°C using 20units of T4 RNA ligase 1 (NEB) in the presence of 10% PEG 8000 to favour intramolecular ligations. Ligated RNAs were then purified using phenol/chloroform extraction followed by ethanol precipitation and then digested using 10U RNase R from Epicentre as described in the RNase R treatment paragraph. Between 24h and 72h following siRNA transfection, 0.036µg of *in vitro* synthesized circular RNAs were transfected in WI38 RAF1-ER cells, using Dharmafect 4 (Dharmacon) according to the manufacturer’s recommendations.

### *In vitro* RNA synthesis for pull-down experiments

10µg of plasmid (Table S5) were digested overnight using PvuII for scramble and SmaI for circANRIL. Linearised plasmids were used as template for *in vitro* transcription using T7 RNA polymerase (Promega) and biotin RNA labeling mix (Sigma Aldrich) following the manufacturer’s recommendations for 2h at 37°C. *In vitro* transcribed RNA were incubated for 1h at 37°C after addition of 10units DNase I and 10units DNase Zero (Epicentre). The RNA were then ethanol precipitated and resuspended in water and dephosphorylated using 2units Alkaline Phosphatase (FastAP, Thermo Scientific) for 30min at 37°C following the manufacturer’s recommendations. De-phosphorylated RNAs were purified by phenol/chloroform extraction followed by ethanol precipitation, re-phosphorylated using 10units T4 PNK (Promega) for 30min at 37°C and ethanol precipitated. Around 500ng of RNA were ligated overnight at 16°C using 20units of T4 RNA ligase 1 (NEB) in the presence of 10% PEG 8000 to favour intramolecular ligations. Ligated RNAs were then purified using phenol/chloroform extraction followed by ethanol precipitation. For four of the pull-down experiments (out of eight independent experiments), we used the MasterPure RNA purification kit from Epicentre instead of phenol/chloroform extraction to purify RNA after the dephosphorylation and ligation steps.

### Biotinylated RNA pull-down

30µL of HeLa nuclear extract (purchased from Computer Cell Culture Center, prepared according to the classical Dignam protocol) were diluted 10 times in binding buffer (20mM Tris pH 8.0, 100mM NaCl, 5mM MgCl2, 0.4% NP40) and supplemented with protease inhibitors (Roche), 60µg of yeast tRNA (Invitrogen) and 8U of DNase I (Epicentre). These diluted nuclear extracts were pre-cleared using 25µL of previously blocked Streptavidin Sepharose beads (GE Healthcare) for 2h at 4°C. Blocking was achieved by incubating the beads with 1mg.mL^-1^ of Ultrapure BSA and 0.5mg.mL^-1^ of salmon sperm DNA (Invitrogen) overnight at 4°C. The precleared nuclear extracts were recovered, supplemented with 2µL Riboguard (Epicentre) and incubated for 30min at room temperature with 500ng of *in vitro* synthesized RNAs. 10µL of blocked beads were then added and incubation was pursued for another 30min at room temperature. After 4 washes in binding buffer, beads were resuspended in 20µL binding buffer and 10µL 4X Laemmli sample buffer (Biorad) supplemented with b-mercaptoethanol. Samples were boiled for 5min at 95°C before being loaded on a 3-8% NuPAGE Tris-Acetate gel (Invitrogen). Western blots were quantified using ImageJ. Linearity of the EZH2 and SUZ12 western blots was controlled by serial dilutions of an input.

### RNA FISH

The protocol used for RNA FISH experiments has been described elsewhere (46) except that probes were denatured for 7 minutes at 75°C and additionally pre-incubated 10 min at 37°C. Red-dUTP (02N34-050, Abbot Molecular) labeled probes were generated by nick translation using the WI2-48403-G248P84444H2 fosmid. All images were taken on a fluorescence microscope DM5000 B Leica with a Retiga R3 camera controlled by the Metamorph 7.7 software using x40 objective. Different wavelength probes were used (DAPI (360nm, 470nm) and Texas Red (596nm, 612nm)). Images were quantified using Columbus, the integrated software of the Operetta automated high-content screening microscope (PerkinElmer).

### Accession numbers

RNA Seq data are available at GEO (**<u>GSE 143957</u>**).

## Abbreviations

ncRNAs: non-coding RNAs;
OIS: oncogene-induced senescence;
*ANRIL*: antisense non- coding RNA in the *INK4* locus;
CDK: Cyclin Dependant Kinase;
CAD: coronary artery diseases;
circRNAs: circular RNAs;
SAHF: senescence associated heterochromatin foci;
SASP;: senescence associated secretory phenotype;
ER: estrogen receptor;
hTERT: human telomerase reverse transcriptase;
4-HT: 4-hydroxy-tamoxifen;
RNA pol II: RNA polymearse II;
EdU: 5-ethynyl-2’-deoxyuridine;
EU: 5-ethynyl uridine;
RT-qPCR: reverse transcription-quantitative PCR;
ChIP: chromatin immunoprecipitation;
FISH: fluorescence *in situ* hybridization;
RNA-seq: RNA sequencing;

## Acknowledgments

This work was supported by grants from the INCa (PLBio program), the Fondation ARC (programme ARC), the Fondation Toulouse Cancer Santé and the Ligue Nationale Contre le Cancer (LNCC, équipe labélisée) to DT and from the Fondation de France to LM. EN is employed by INSERM. LM was supported by fellowships from the LNCC, the Fondation ARC and the Fondation de France. MA was supported by a fellowship from the Fondation pour la Recherche Médicale (FRM), SL was supported by studentships from the Fondation ARC and JO by a studentship from the French Ministry of Research. We thank Malek Djabali and Jacques Côté for constructive discussions throughout the study, Céline Vallot and Claire Rougeulle for sharing their expertise on nuclear RNA FISH.

---

**Supplementary Figure 1:**
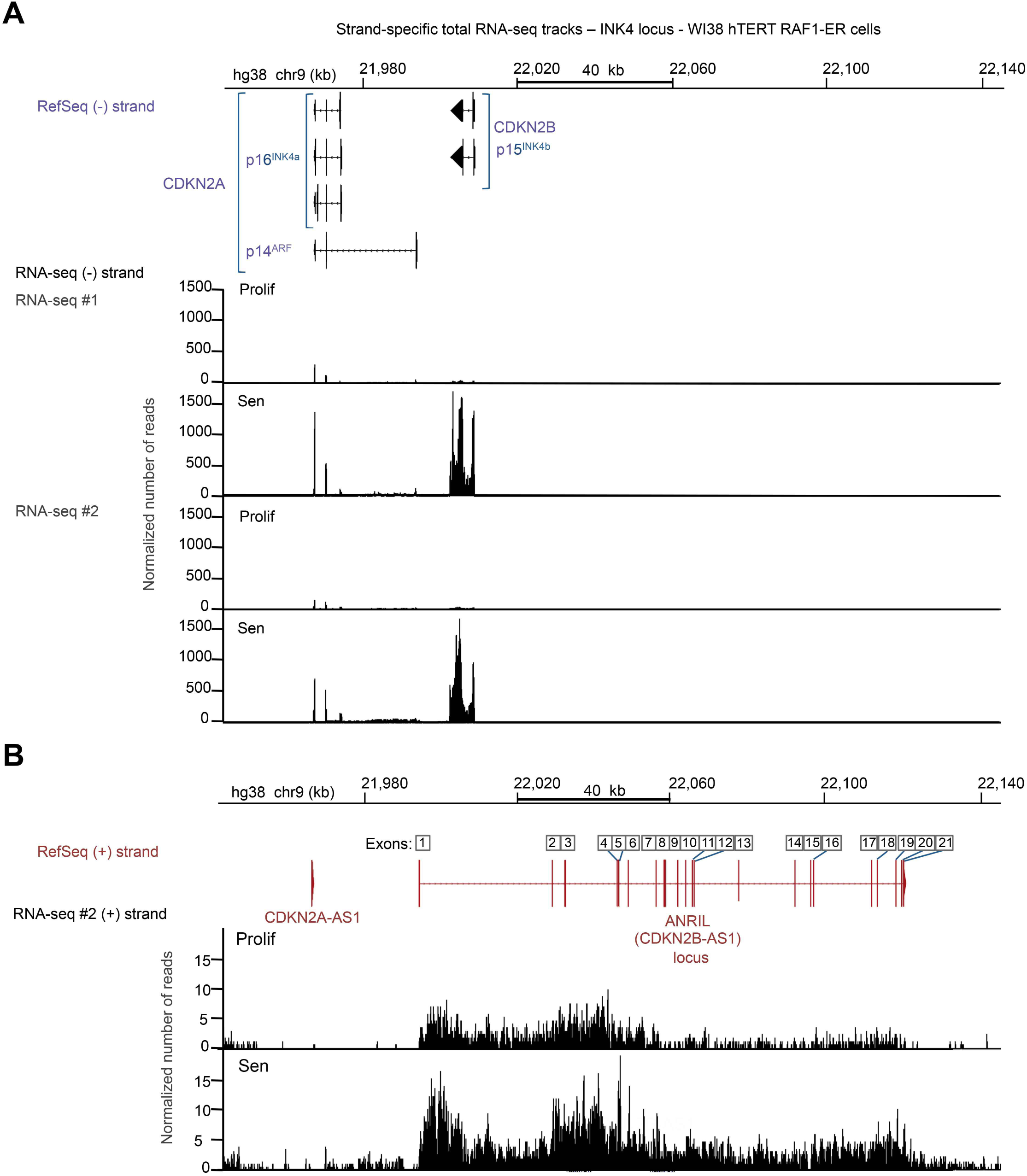
Strand-specific total RNA-seq from proliferating and senescent Wl38 hTERT RAF1-ER cells. RefSeq Genes (hg38) and tracks from strand-specific total RNA-seq in Wl38 hTERT RAF1-ER cells induced (Sen) or not (Prolif) in senescence are shown at the *INK4* locus for the minus DNA strand of two independent replicates of RNA-seq (A) and for the plus strand of the second replicate of RNA-seq (B). Because *ANRIL* has many transcript variants, a concatenated transcript of *ANRIL* (ANRIL locus), which comprises all the exons present in all the transcript variants described in the reference genome (RefSeq, hg38), is shown.

**Supplementary Figure 2:**
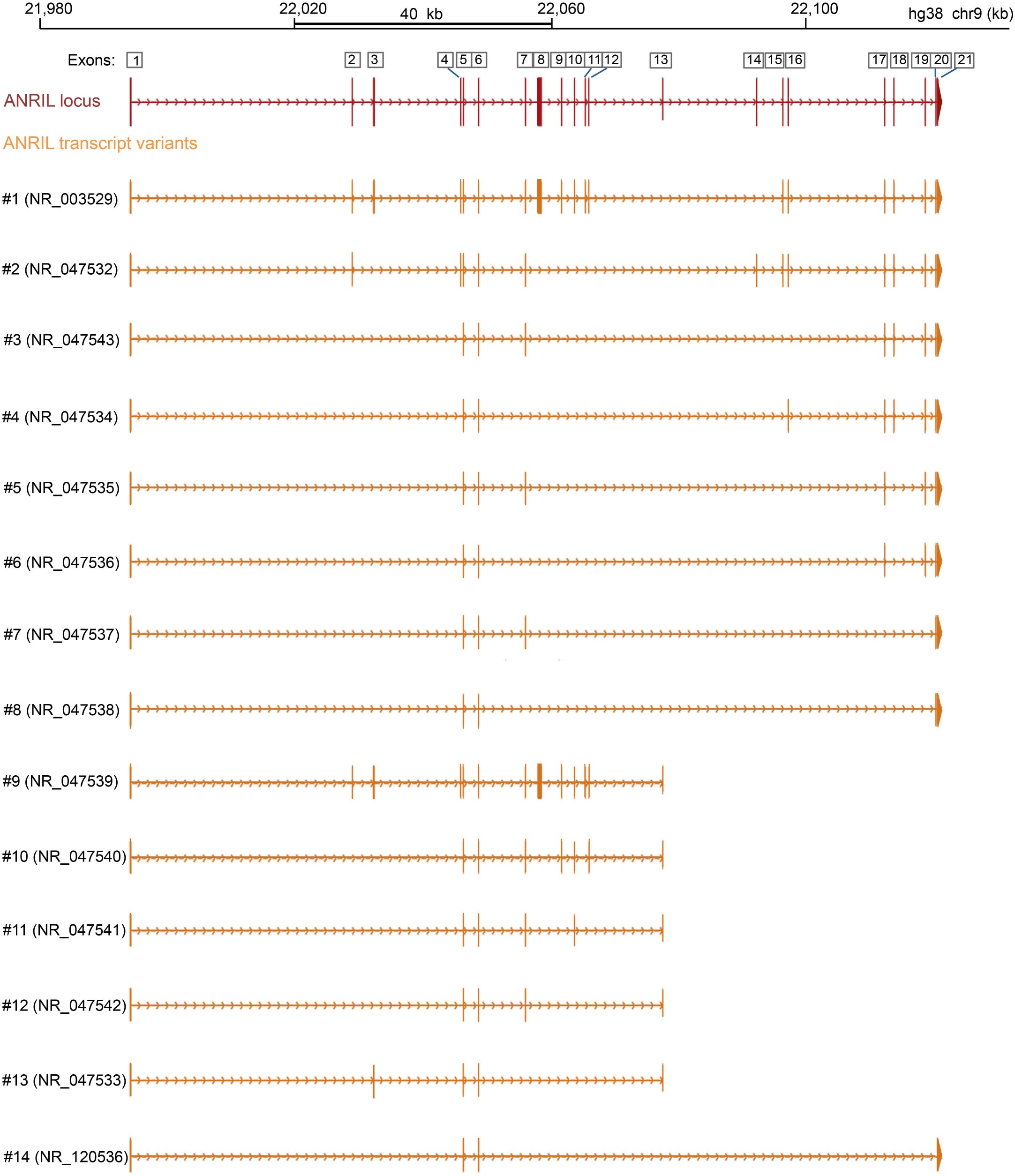
Schematization of *ANRIL* locus with the numbering of all exons present in the various transcript variants of *ANRIL* that are annotated in the reference genome (RefSeq hg38).

**Supplementary Figure 3:**
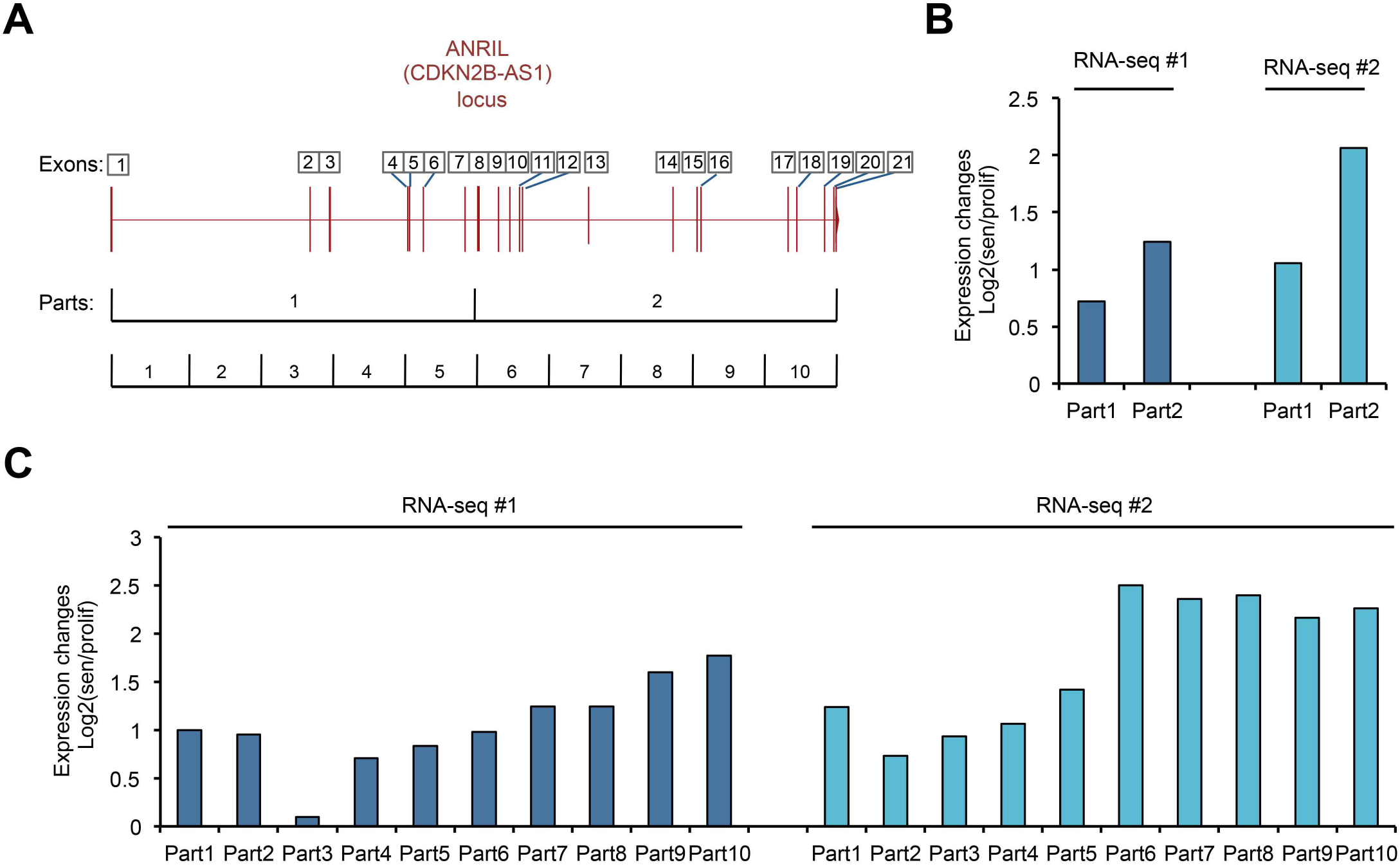
*ANRIL* transcript increases more in senescence in its second half than in its first half. The *ANRIL* gene locus was divided in two (B) or in ten (C) equal parts, as shown in panel A). For each of these regions, the mean per base of the normalized number of aligned reads was computed. The fold expression changes (Log2 ratio sen/prolif) in senescence of *ANRIL* expression was calculated in each of these parts in the two RNA-seq replicates.

**Supplementary Figure 4:**
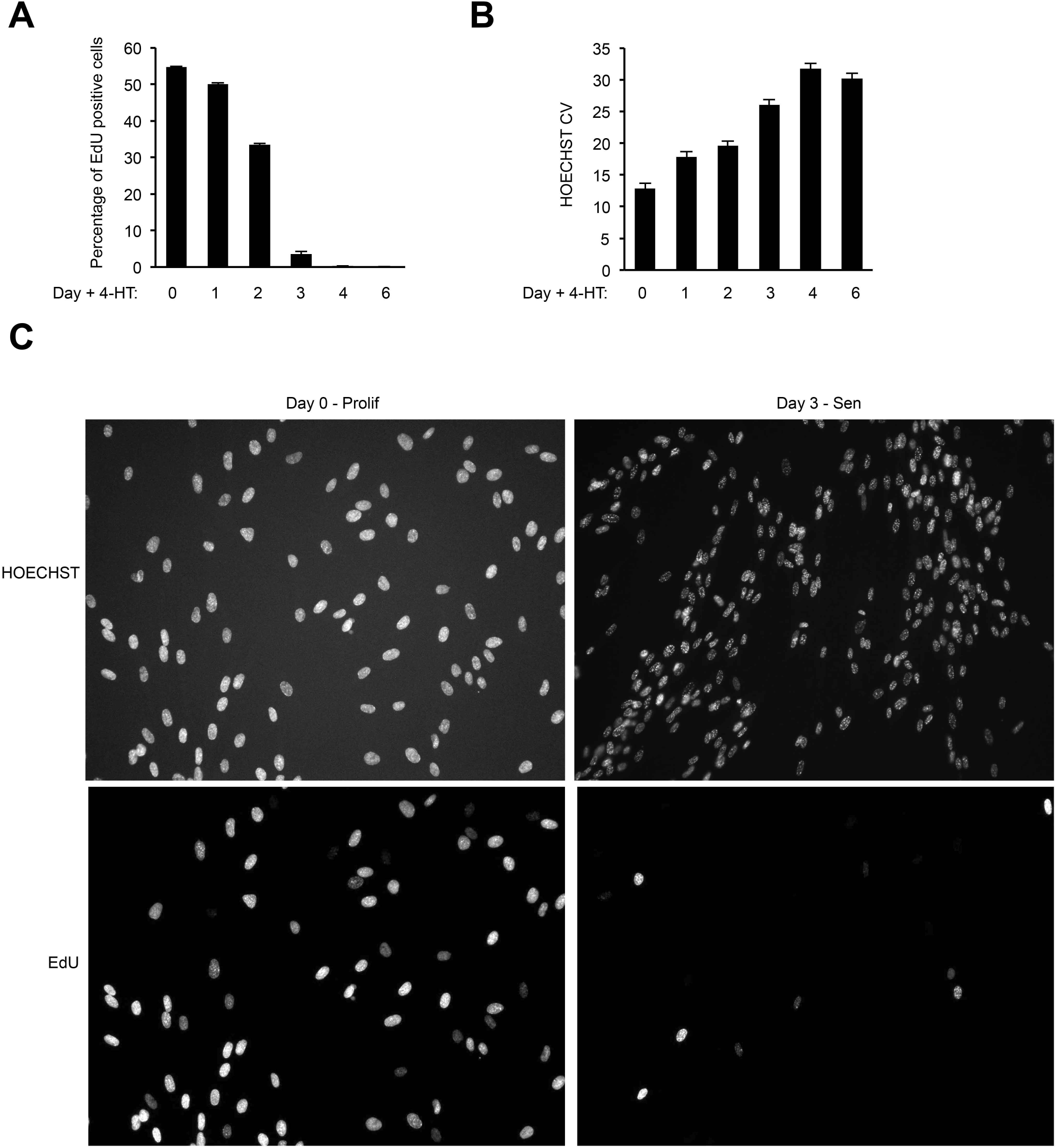
Kinetics of senescence induction in Wl38 hTERT RAF1-ER. **A)** Proliferative Wl38 hTERT RAF1-ER cells were treated with 4-hydroxytamoxifen (4-HT) for the indicated times in a 96 wells plate in duplicates. Cells were treated with EdU for 24 hours before being fixed, stained with Hoechst and analyzed using the Operetta device (Perkin Elmer). At least 3900 cells were analyzed in each sample. The percentage of EdU positive cells was calculated for each time point. A representative experiment out of two is shown (means and standard deviations of the duplicates). **B)** Same as in (A), except that Hoechst CV (Coefficient of Variation, representing SAHF formation) was calculated in each cell. A representative experiment out of two is shown (means and standard deviations of the duplicates). **C)** Representative images obtained by the Operetta microscopy device of proliferative (day 0) and senescent (day3) Wl38 hTERT RAF1-ER.

**Supplementary Figure 5:**
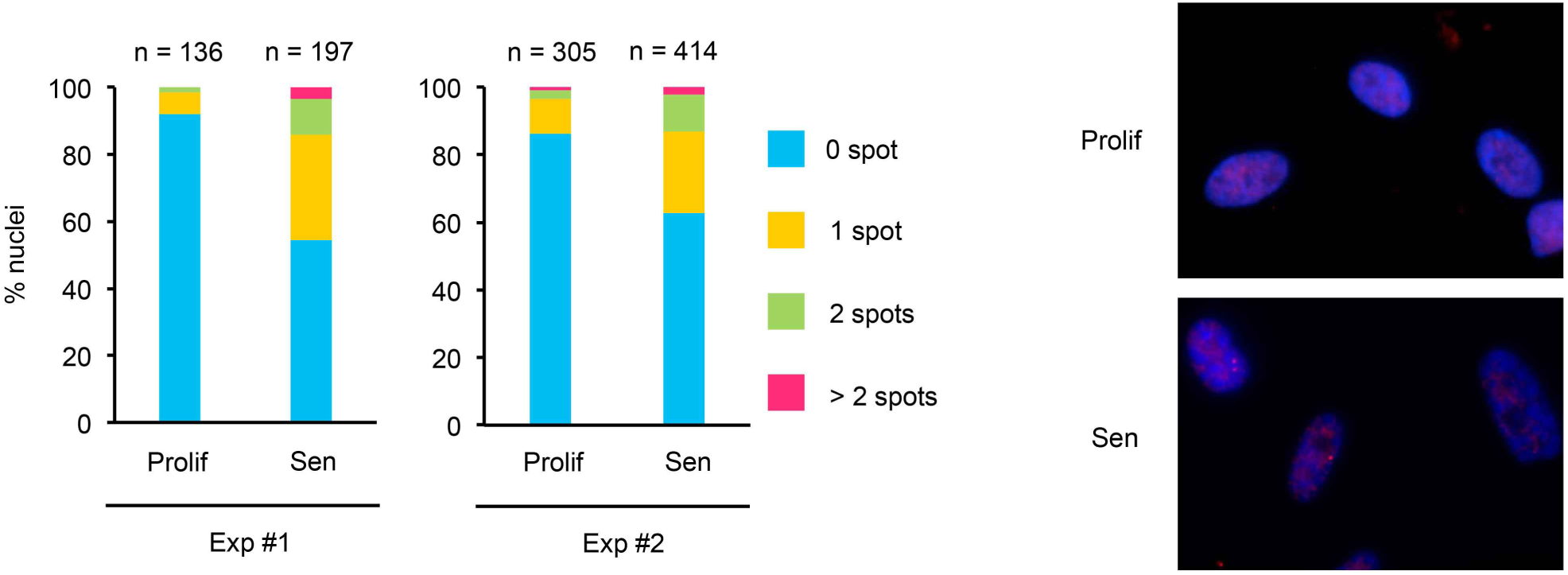
Increase of *ANRIL* expression at the individual cell level during RAF1-induced senescence. Total *ANRIL* RNA FISH experiments in proliferative and senescent Wl38 hTERT RAF1-ER cells. The probe used in these experiments is generated from a fosmid covering the entire genomic region of *ANRIL*. The number of spots were counted in each nucleus from two independent experiments (Exp #1 and Exp #2). The numbers of analyzed cells are shown on top of the graphs. Representative images are shown in the right panel. Note that the protocol we used for the RNA FISH experiments is designed to label nuclear RNAs and not cytoplasmic RNAs.

**Supplementary Figure 6:**
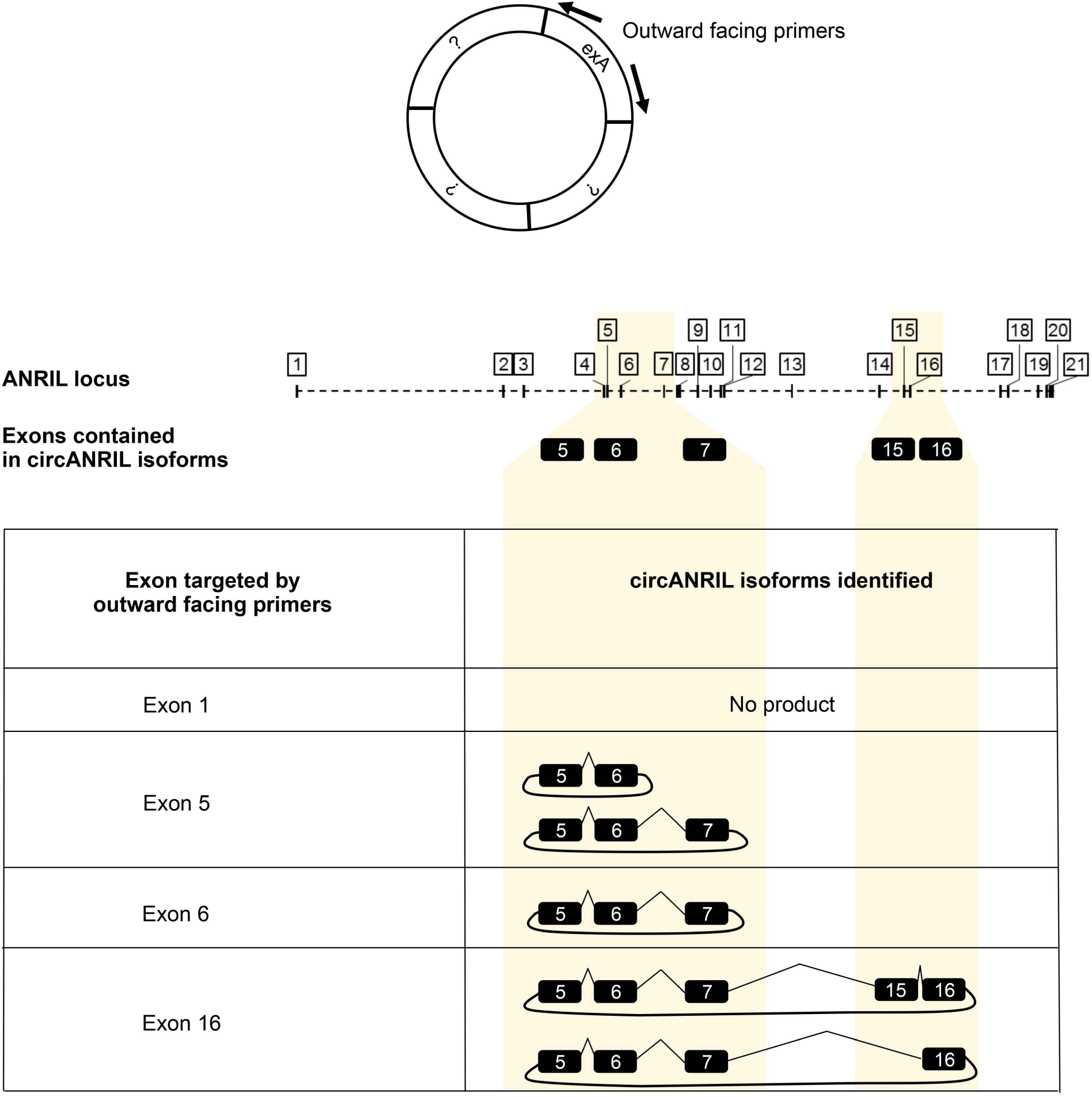
Identification of circular *ANRIL* species. Total RNAs from senescent Wl38 hTERT RAF1-ER cells were used for random primed reverse transcription followed by PCR using outward facing primers (which amplify circular but not linear molecules, as schematized) designed in several different exons of *ANRIL* (exon1; exon 5; exon 6 and exon 16). PCR products were then run on a 1% agarose gel, bands were cut out and extracted from the gel and sequenced with either one of the primers used for the PCR. Sequencing of the PCR products allowed us to identify four different circular *ANRIL* species as schematized.

**Supplementary Figure 7:**
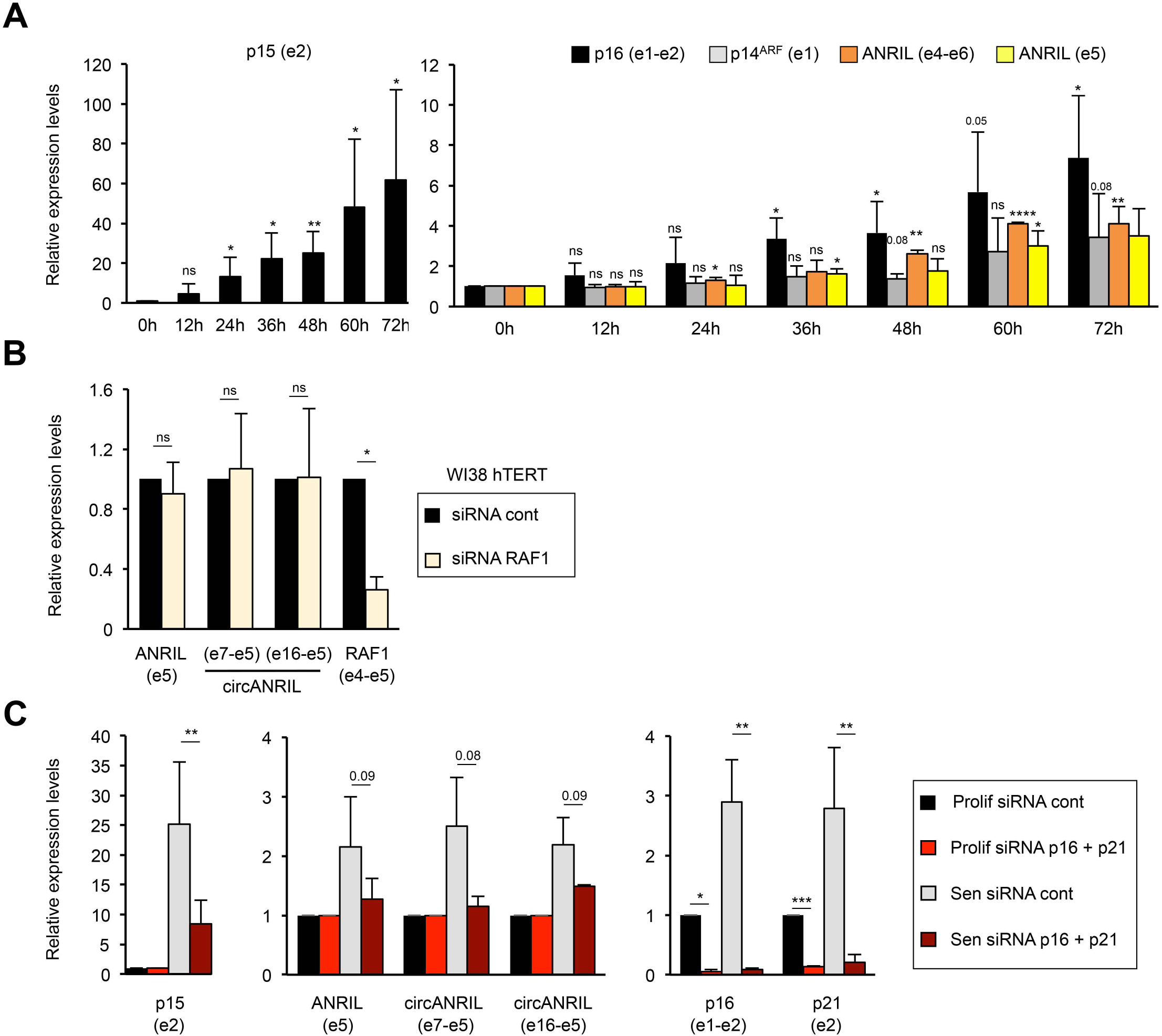
Increase in *ANRIL* and *circANRIL* expression during senescence induced by **the RAF1 oncogene**. **A)** Short kinetics of RAF1-induced senescence. Wl38 hTERT RAF1-ER cells were induced to senescence for the indicated time in hours (h), followed by total RNA extraction. RNA expression was measured by RT-qPCR using the indicated primers. The means and standard deviations from 3 independent experiments are shown (except for ANRIL e5 at the 72h time point, for which only 2 experiments were analyzed), relative to *GAPDH* and normalized to 1 at the time Oh. Two-tailed paired Student’s t-tests were applied on log2 values between time Oh and other times of the kinetics for each primer pair. Significant differences are indicated by an asterisk (*: p value < 0.05, ** to ****: p values < 10-2, 10-3 and 10-4, respectively); number of the p value is indicated when it is between 0.05 and 0.1; ns: not significant. **B)** Depletion of RAF1 in Wl38 hTERT cells does not decrease *ANRIL* and *circANRIL* expression. Wl38 hTERT cells (which do not contain RAF1-ER construct) were transfected with the indicated siRNAs (both from Dharmacon). Following extraction, RNA expression was measured by RT-qPCR using the indicated primers. The means and standard deviations from 3 independent experiments are shown, relative to *GAPDH* and normalized to 1 in siRNA cont cells. Significant differences are indicated as in A). **C)** The increase of *ANRIL* and *circANRIL* expression in RAF1-induced senescence is associated with senescence. Wl38 hTERT RAF1-ER cells were transfected by the indictated siRNAs (control Dharmacon), including p16 and p21 siRNAs, which inhibits senescence induction (Jeanblanc et al., 2012). The next day, cells were induced (Sen) or not (Prolif) in senescence for 3 days. Following extraction, RNA expression was measured by RT-qPCR using the indicated primers. The means and standard deviations from 3 independent experiments are shown, relative to *GAPDH* and normalized to 1 in proliferative cells for p15, ANRIL (e5), circANRIL (e7-e5) and circANRIL (e16-e5) or to 1 in proliferative cells treated with control siRNA for p16 and p21. Depletion of p16 and p21 did not affect the expression of *ANRIL* and *circANRIL* in proliferative cells (data not shown). The induction of *ANRIL* and *circANRIL* in senescent cells is lower in these experiments as compared with experiments shown in Fig. 1D and 3B, probably due to the stress induced by the siRNA transfection. Significant differences are indicated as in A).

**Supplementary Figure 8:**
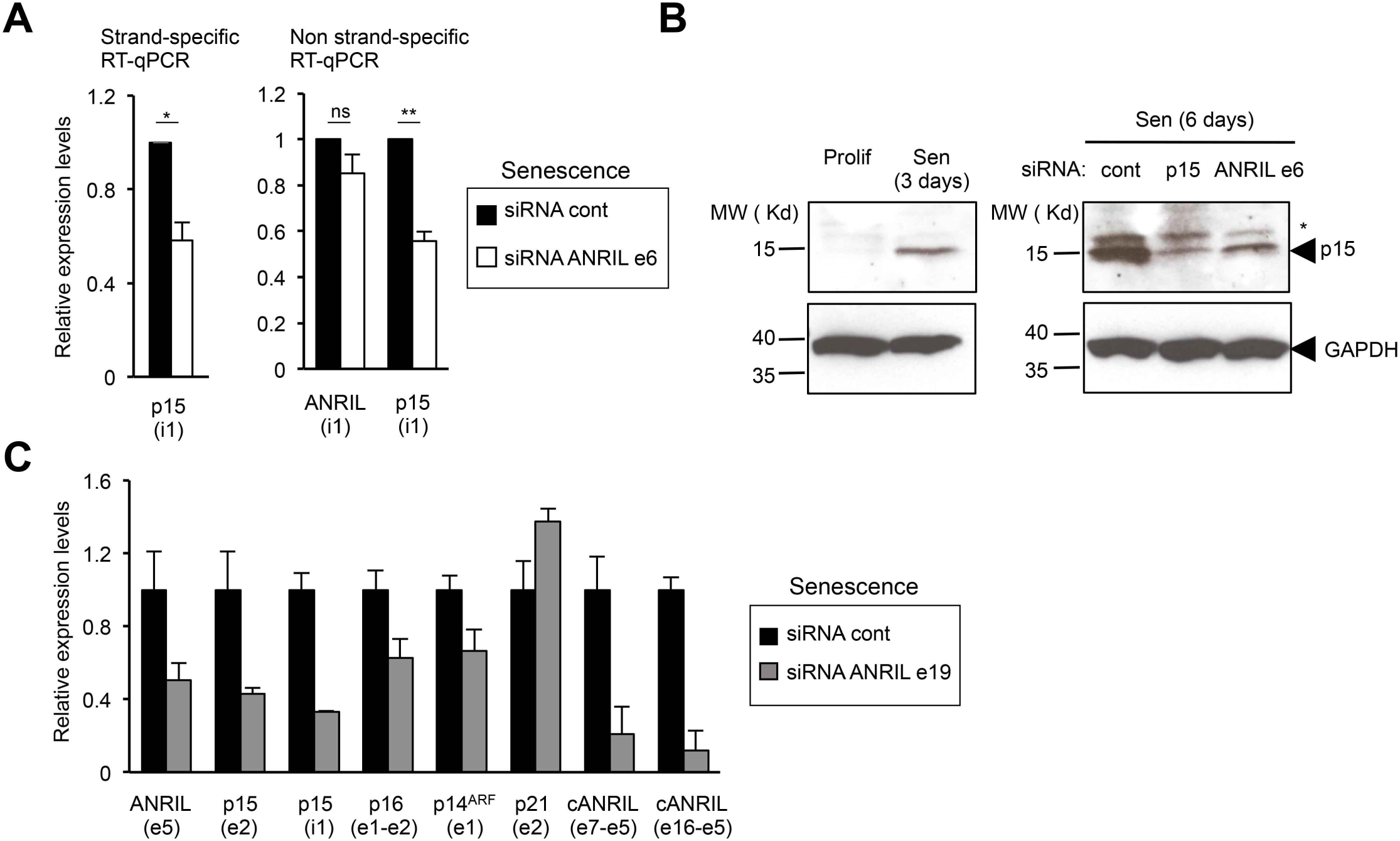
Depletion of total *ANRIL* leads to a decrease of p15 gene expression as well as of p16 and p14^**ARF**^ **gene expression in RAF1-induced senescence**. **A)** Control of the specificity of the detection of p15 pre-mRNA, to which ANRIL is antisense, by random priming reverse transcription. Total RNA from senescent Wl38 hTERT RAF1-ER cells transfected with siRNA targeting the exon 6 of *ANRIL* or control (pool) siRNAs were extracted. Left panel: RNAs were used for strand-specific reverse transcription using either p15 (i1) or GAPDH (e9) reverse primers. The cDNAs were then used for qPCR using p15 (i1) or GAPDH (e9) primers. *pt5* (i1) levels are expressed relative to *GAPDH* (e9) levels and normalized to 1 in siRNA control-treated cells. The means and standard deviations from 3 independent experiments are shown. Significant differences are indicated with asterisks (*: p value < 0.05, ** to **** p values< 10-^2^, 10-^3^ and 10-^4^ respectively, two-sided paired Student’s t-test on log2 values); ns: not significant. Right panel: RNAs were used for non strand-specific reverse transcription using random primers and the expression of *ANRIL* or *pt5* was measured using the indicated primers. ANRIL (i1) primers used for qPCR were designed outside of the region antisense to *pt5* gene to avoid monitoring *pt5* expression. The means and standard deviations from 3 independent experiments are shown. Significant differences are indicated as in the left panel. Note that *pt* 5 pre-mRNA expression (measured in intron 1 (i1), which is on the antisense DNA strand to *ANRIL* intron 1 sequence, see Fig. S1) is decreased in the siRNA ANRIL ex6 sample, as measured by strand-specific or non strand-specific reverse transcription. The expression of *ANRIL* pre-mRNA (measured in intron 1 (i1) in the non-complementary region to *pt* 5) is unaffected by the siRNAs against ANRIL e6. Moreover, the expression level of *pt* 5 measured in its intron 1 is around 16 times higher than the expression level of *ANRIL* measured in its intron 1 (data not shown). Altogether, this indicates that when we monitor *pt5* intron 1 expression by random priming, in the right panel, in C), in Fig. 5, S9A and S10A, the results are unaffected by the levels of *ANRIL* intron 1. **B)** Depletion of *ANRIL* decreases p15 protein expression. Proliferative Wl38 hTERT RAF1-ER cells, senescent cells (harvested 3 days following 4-HT addition) or senescent cells transfected by the indicated siRNAs for 72 hours (thus harvested 6 days following senescence induction) were subjected to whole cell protein extraction. p15 and GAP DH expression were then monitored by western blot. One representative experiment out of 2 is shown. The band marked by an asterisk is certainly a non- specific band, as it does not decrease upon p15 depletion. **C)** Depletion of *ANRIL* using an siRNA directed against the exon 19 of *ANRIL* also leads to decreased expression of *INK4* genes in RAF1-induced senescence. Total RNAs from senescent Wl38 hTERT RAF1-ER cells transfected with siRNA targeting the exon 19 of *ANRIL* or control (cont3) siRNA were extracted. The expression of the indicated RNAs was measured using the indicated primers. One representative experiments out of two is shown (means and standard deviations from qPCR triplicates).

**Supplementary Figure 9:**
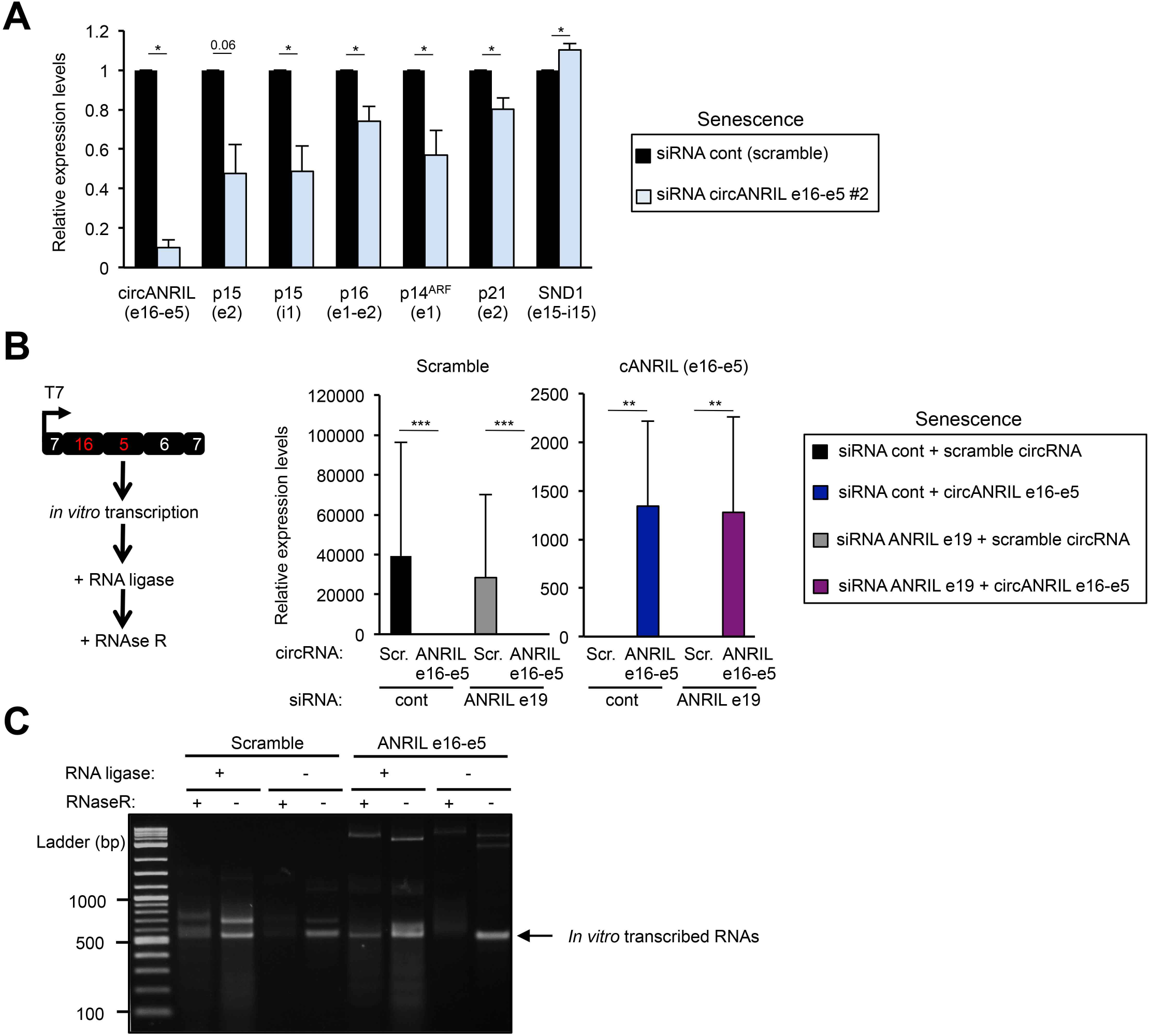
*CircANRIL* e16-e5 participates to *INK4* gene activation in RAF1 -induced senescence. **A)** Senescent Wl38 hTERT RAF1-ER cells were transfected with the indicated siRNAs. siRNA circANRIL e16-e5 #2 targeting circANRIL e16-e5 back-spliced junction is different from the siRNA used in Fig. 5 by 3 nucleotides. As control, a siRNA scramble of this sequence was used. Total RNAs from transfected cells were subjected to RT-qPCR with the indicated primers. Means and standard deviations from 3 independent experiments, calculated relative to *GAPDH* mRNA and normalized to 1 for the control siRNA. Significant differences are indicated with asterisks(*: p value< 0.05, **to**** p values< 10-2, 10-3 and 10-4 respectively, two-sided paired Student’s t-test on log2 values); the number of the p value is indicated when it is between 0.05 and 0.1. **B)** *In vitro* transcription was performed to produce the circANRIL e16-e5 sequence or a scramble (Ser.) sequence (which does not correspond to any genomic sequence and thus not expressed in normal conditions). Linear RNAs produced *in vitro* were incubated with RNA ligase to circularize the RNAs and then with RNAse R to digest linear RNAs, as schematized. These RNAs were transfected in senescent Wl38 hTERT RAF1-ER cells with siRNAs against the exon 19 of ANRIL (which do not target the *in vitro* circANRIL e16-e5 product) or control siRNAs (cont3). After RNA purification, RT-qPCR were performed using the indicated primers showing the efficient overexpression of circANRIL e16-e5 junction (calculated relative to 1 for siRNA control - scramble circRNA) and of its Scramble RNAs (calculated relative to 1 for siRNA control - circANRIL e16-e5) in the experiments shown in Fig. 5D. qPCR specific for these RNA products also amplified concatemers formed during the RNA ligation reaction, in the last cycles of the qPCR (data non shown). Significant differences are indicated as in A). **C)** Same as B) except that *in vitro* transcribed RNAs were treated or not with RNA ligase and Rnase R and run on an agarose gel to check their size (expected size: 512 nucleotides) and the ligation process, which induces circularization and protects the products from RNAse R. The same protocol was used to produce the RNAs used in the RNA pull down experiments shown in Fig. 6B, except that a biotin RNA labelling mix containing biotin-16-UTP was used for the *in vitro* transcription step.

**Supplementary Figure 10:**
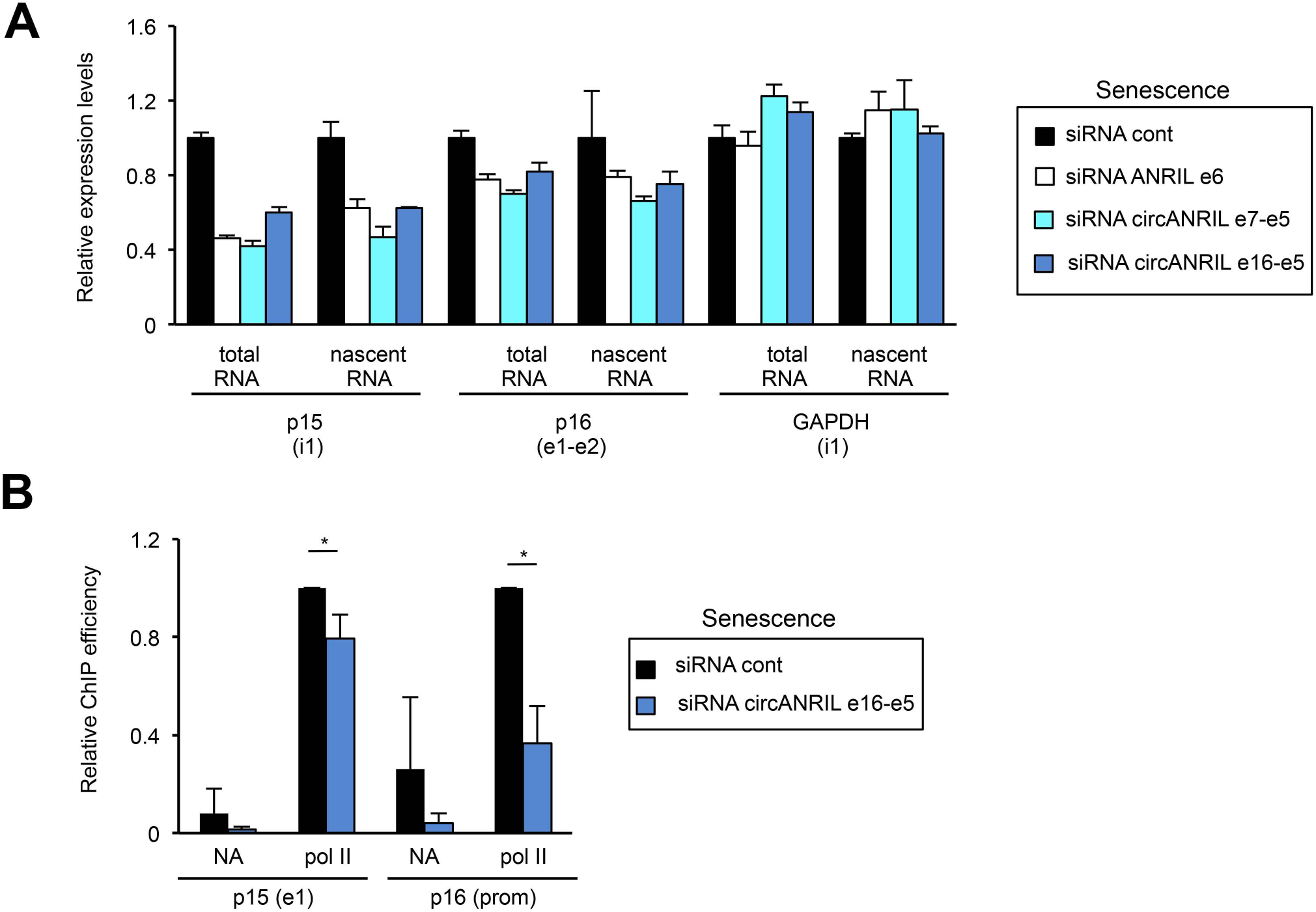
*CircANRIL* e7-e5 and e16-e5 isoforms favor *INK4* gene transcription in RAF1-induced senescence. **A)** *CircANRIL* isoforms favor *p15* transcription as assessed by nascent RNA capture experiments. Senescent Wl38 hTERT RAF1-ER cells were transfected with the indicated siRNAs (siRNA cont3 for control). 72 hours later, cells were incubated with EU to label nascent RNAs for 10 or 30 min. Total RNAs were extracted and subjected or not to nascent RNA capture. *p15* or *GAPDH* (as control) measured both in their intron 1 and *p16* were analyzed, calculated relative to *GAPDH* mRNA (measured in exon 9) and normalized to 1 for the control siRNA. One representative experiment out of 2 is shown. The means and standard deviations from the qPCR triplicates are shown. **B)** *CircANRIL* e16-e5 depletion inhibits RNA pol II recruitment to the *p15* and *p16* promoters. Senescent Wl38 hTERT RAF1-ER cells were transfected with the circANRIL e16-e5 siRNA. Three days later, cells were harvested and subjected to a ChlP analysis using RNA pol II antibody or no antibody (NA) as a control. The amount of p15 e1, p16 e1 or GAP DH e1 (used as a positive control) sequences were quantified by qPCR. The percentage of input was calculated and the amount of *p15* and *p16* promoters were calculated relative to 1 for the control siRNA sample following standardization with GAPDH e1 sequences. The means and standard deviations from 3 or 4 independent experiments are shown. Significant differences are indicated by an asterisk (p value < 0.05, two-sided paired Student’s t-test on log2 values).

**Supplementary Figure 11:**
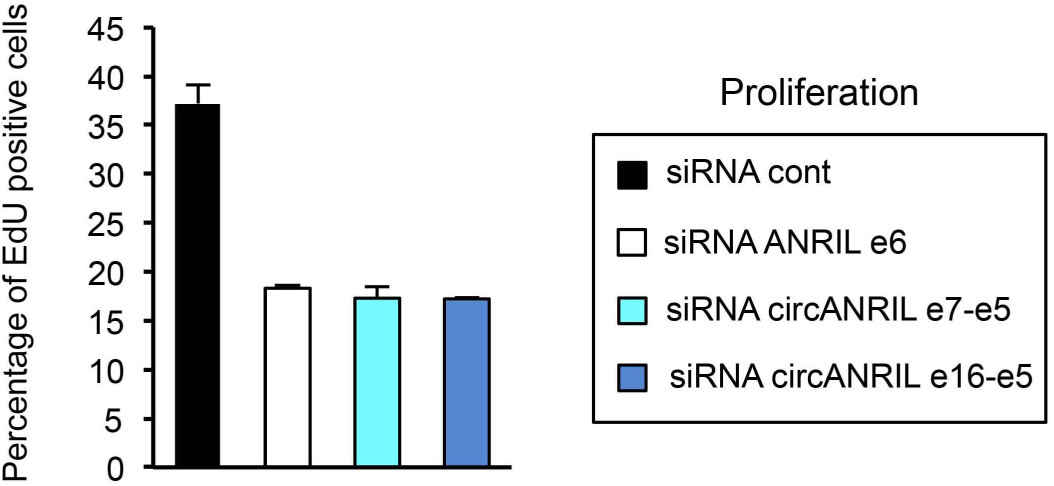
Depletion of *circANRIL* isoforms in proliferative cells leads to proliferation arrest. Proliferative Wl38 hTERT RAF1-ER cells were transfected in a 96 wells plate in duplicates using the indicated siRNAs (siRNA cont3 for control). 72 hours later, cells were treated with EdU for 24h, fixed, and analyzed using the Operetta device (Perkin Elmer). At least 5500 cells were analyzed in each well. One representative experiment out of two is shown (means and standard deviations from the 2 duplicates).

## Supplementary Table Legends for Tables S1-S4

**Supplementary Table 1: Analysis of spliced junctions of *ANRIL* from RNA-seq datasets in proliferation and senescence, including circular junctions**

All the back-spliced junctions found in the RNA-seq datasets in proliferation and senescence are shown for the *ANRIL* locus. For regular spliced junctions, only the splicing events with a frequency of 1 read or more in at least one sample (proliferation or senescence) in both RNA-seq replicates are shown. e: exon, i: intron, Exact match: corresponds to the exact boundaries of exons, intg: intergenic

**Supplementary Table 2: List of regulated genes by *circANRIL e16-e5***

Differential analysis of gene expression using DESeq2 package was performed on RNA-seq experiments following depletion of *circANRIL* e16-e5 in senescence. Genes of interest were selected when |log2FoldChange| was higher than 1 and adjusted p-value (padj) lower than 0.05.

**Supplementary Table 3: Gene ontology analysis on the list of up-regulated genes upon *circANRIL* e16-e5 depletion**

Gene ontology analysis was performed using GeneCodis. Only pathways enriched with 5 or more genes are shown.

**Supplementary Table 4: Gene ontology analysis on the list of down-regulated genes upon *circANRIL* e16-e5 depletion**

Gene ontology analysis was performed using GeneCodis. Only pathways enriched with 5 or more genes are shown.

**Supplementary Table 5: List of sequences for primers, siRNA sand constructs for *in vitro* transcription**

F: Forward; R: Reverse; S: Sense; AS: Antisense

Primers for GAPDH (e9), p15 (e2), p16 (e1-e2), p14^ARF^ (e1), p21 (e2) have previously been described in Lazorthes et al (1), for ANRIL (e4-e6) in Liu et al (2), for p16 (prom, -915 nt) (3) in and for U2 in Vilborg et al (4).

All primers and siRNAs (with UU overhangs and 5’ phosphate of the antisense strand, HPLC purification) were ordered at Eurogentec, except for p16/CDKN2A siRNA (Custom ON Target, CCAACGCACCGAATAGTTA (5)), p21/CDKN1A siRNA (siGENOME SMARTpool, M-003471-00), RAF1 siRNA (siGENOME SMARTpool, M- 003601-02) and Dharmacon control (ON TARGETplus Non-targeting Pool, D-001810-10), which were ordered at Dharmacon.

For exon and intron numbering in *ANRIL*, the *ANRIL* locus (Fig. S2) was used; for other transcripts, the transcript variant #1 was used.

### Sequences of primers used for qRT-PCR experiments

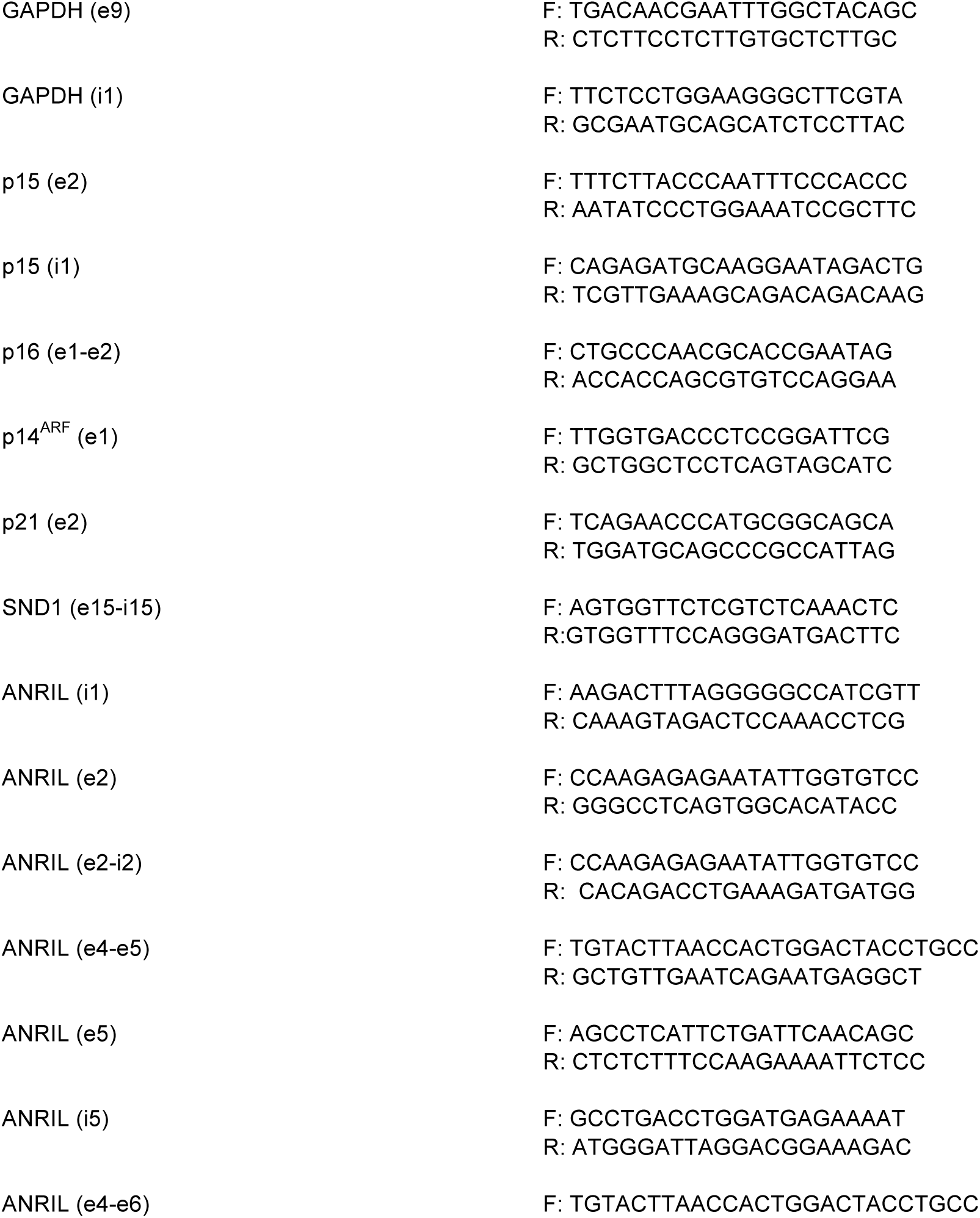
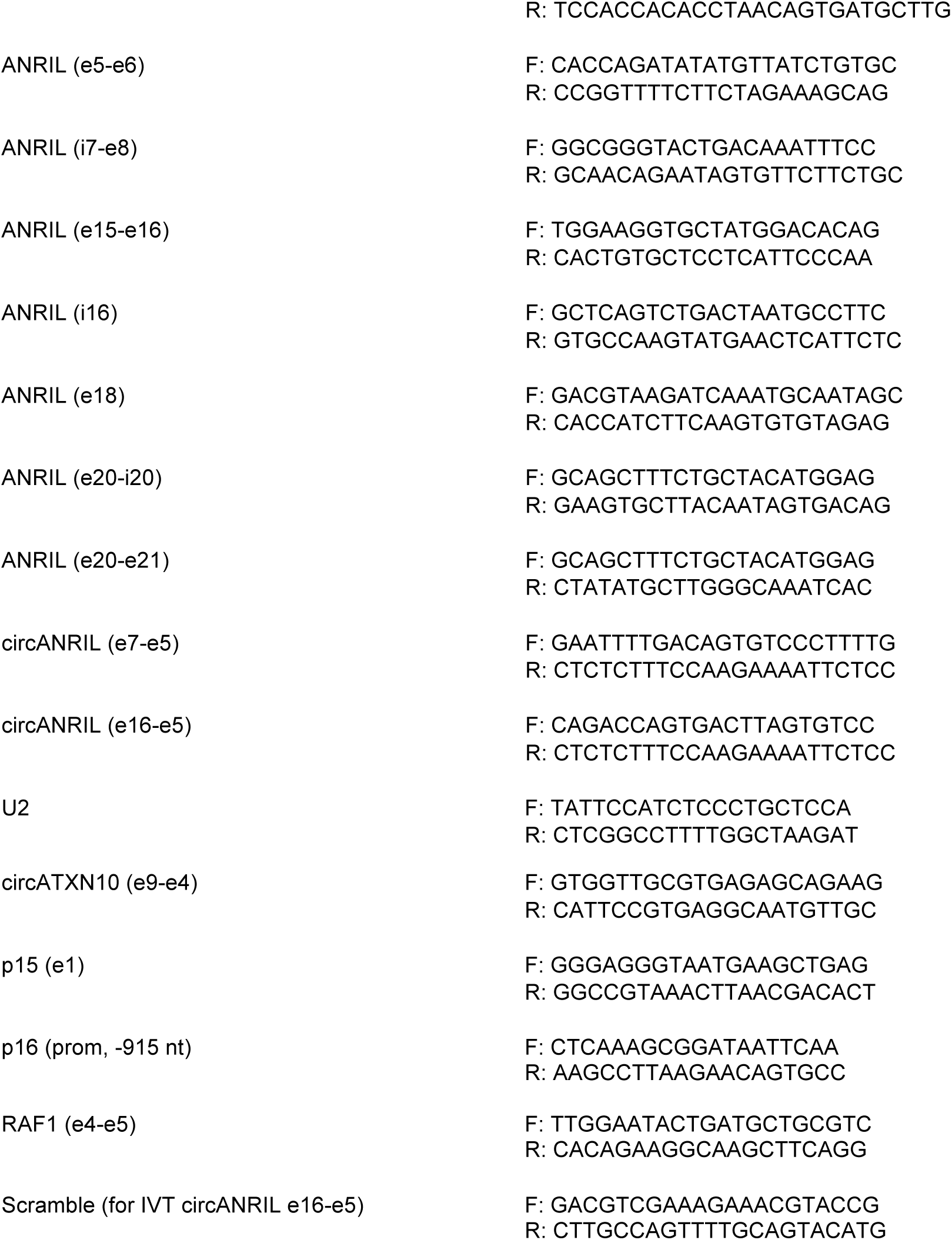

### Sequences of outward facing primers used for PCR

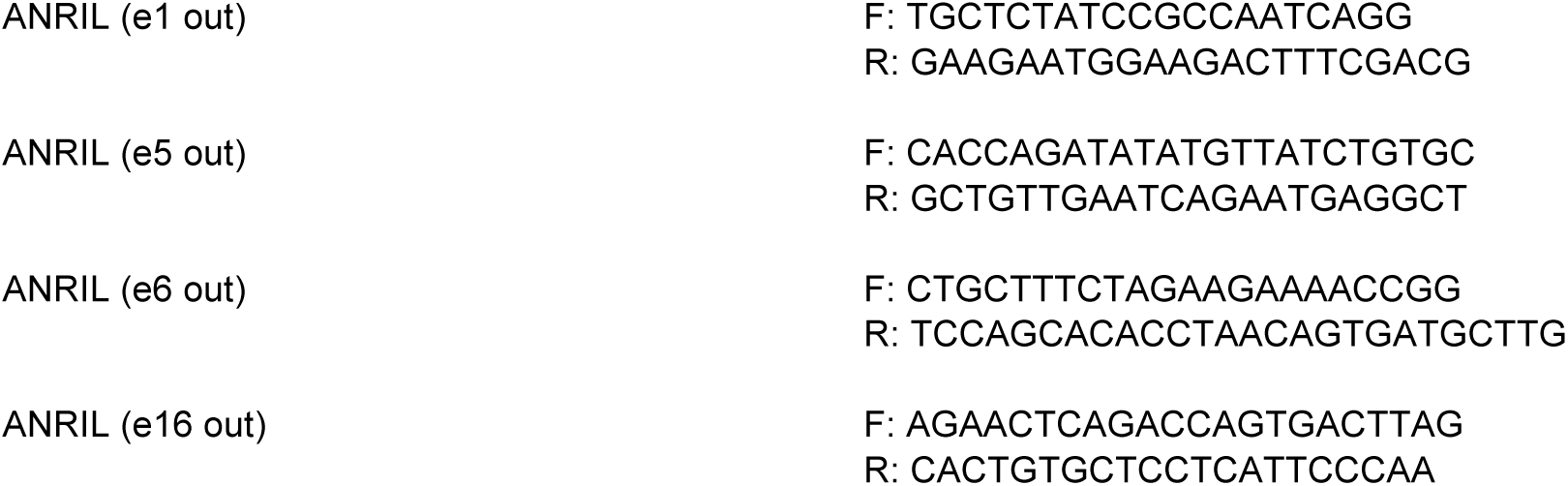

### Sequences of siRNA used

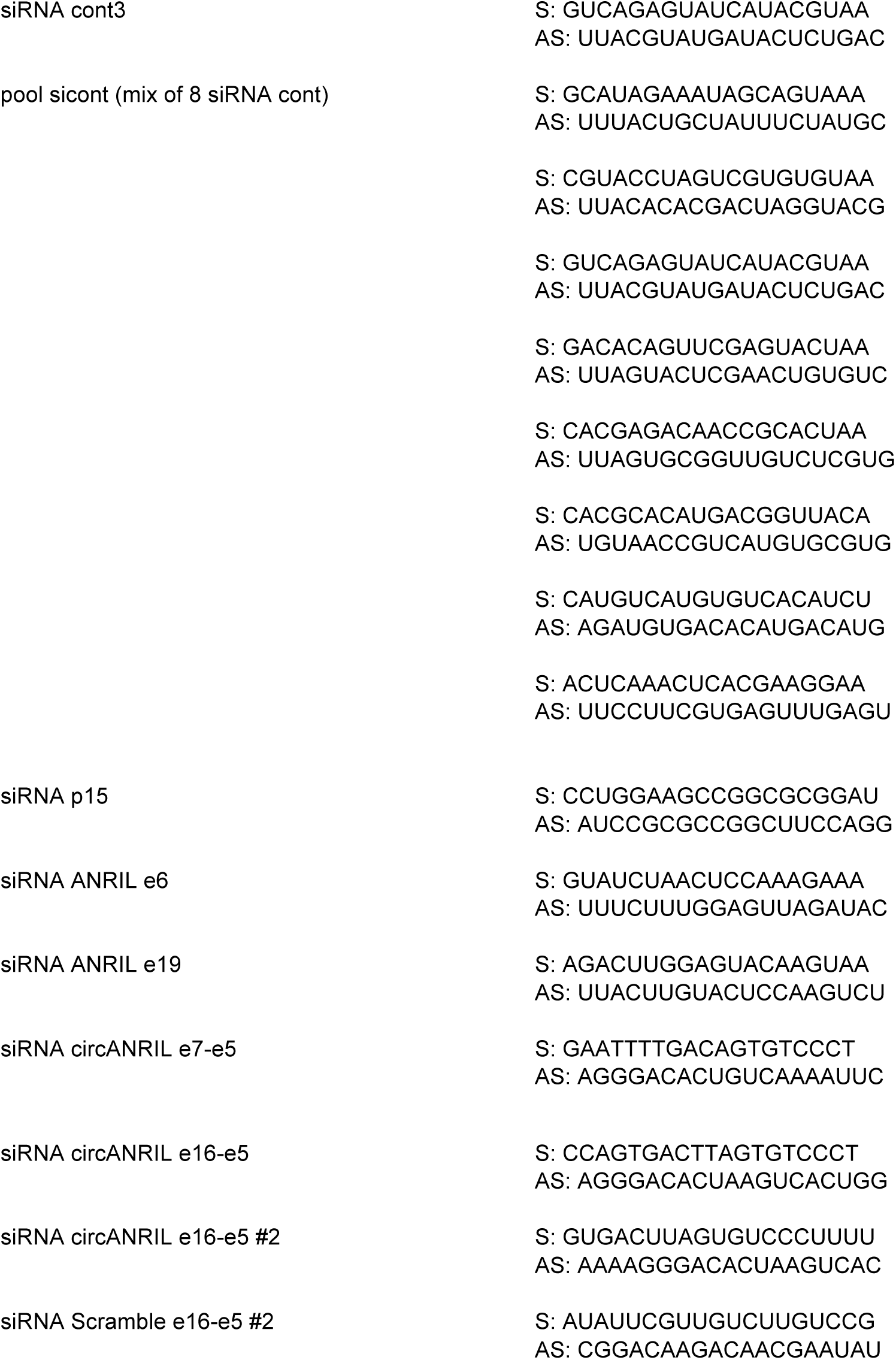

### Sequences of circANRIL e16-e5 and scramble e16-e5 cloned into pUC57 for *in vitro* transcription

IVT Scramble e16-e5

TAATACGACTCACTATAGCGTTCTAAGGAAGGGGCTTGAGAATTCAGGGTTGAGTCATCATTGTCGATTTGCA AGATCTGCTGCCTTTGTAACGTACCATATGAGGTGTTGCAAGCTAGGGTACACCCACATCGTCCCACCTGGC ACGCTACTCTCTGCGGAGAACTAGACGAAATGGAGAGGGAAATTCAAGAAACGAAGACGTCGAAAGAAACGT ACCGGAATTCTAGATTGATCCGCTTCTTGAGCACGGACCCGGTATCGCATGCGAATCGTCAAGCAGGGTACA TAGGATACGGACTCGTATCAAAACGATTTTCAGAAACTGACGCGGTAATGAACATGTACTGCAAAACTGGCAA GCAATGTAACTTTTCCGTCTTAGCTCGCTAAGTCACAAGGGCAGAGAAATACGTTGGACTATATTAAATTGAA CACGCGAGGCAACTGTAACAATTATTATTATTATGTTCTCCAAGTTTATTGCGAGATGGATATATTAGGTAAAG AAAACCGGGAGGGTTTTCGGGGTGCAGCTG

IVT circANRILe16-e5

TAATACGACTCACTATAGCGGATAGAGAGAATTTTGACAGGTGGAGAACTTCAGTAGAGGAAGTGGCAGGAATTTGGGAATGAGGAGCACAGTGATTAAACTGGGGCCATTCATATGAGAGTTTAAGAACTCAGACCAGTGACTTAGTGTCCCTTTTGATGAGAAGAATAAGCCTCATTCTGATTCAACAGCAGAGATCAAAGAAAAGACTTCTGTTTTCTGGCCACCAGATATATGTTATCTGTGCTTAAAGAATTGAAAAACACACATCAAAGGAGAATTTTCTTGGAAAGAGAGGGTTCAAGCATCACTGTTAGGTGTGCTGGAATCCTTTCCCGAGTCAGTACTGCTTTCTAGAAGAAAACCGGGGAGATCTATTTGGAATGTATCTAACTCCAAAGAAACCATCAGAGGTAACAGTAGAGACGGGGTTTCACCATGTTGGCCAGACTGGTCTTGAACTCTCGACCTCGTGATTCGCCCGCCTCGGCCTCCCAAAGTGCTGGG ATTACAGGTGTGAGACACCACACCCGGG

